# Awakening of the RuMP cycle for partial methylotrophy in the thermophile *Parageobacillus thermoglucosidasius*

**DOI:** 10.1101/2024.10.31.621308

**Authors:** Miguel Paredes-Barrada, Annemieke Mathissen, Roland A. van der Molen, Pablo J. Jiménez-Huesa, Machiel Eduardo Polano, Stefano Donati, Miriam Abele, Christina Ludwig, Richard van Kranenburg, Nico J. Claassens

## Abstract

Given sustainability and scalability concerns of using sugar feedstocks for microbial bioproduction of bulk chemicals, widening the feedstock range for microbial cell factories is of high interest. Methanol is a one-carbon alcohol that stands out as an alternative feedstock for the bioproduction of chemicals, as it is electron-rich, water-miscible and can be produced from several renewable resources. Bioconversion of methanol into products under thermophilic conditions (>50°C) could be highly advantageous for industrial biotechnology. Although progress is being made with natural, thermophilic methylotrophic microorganisms, they are not yet optimal for bioproduction and establishing alternative thermophilic methylotrophic bioproduction platforms can widen possibilities. Hence, we set out to implement synthetic methanol assimilation in the emerging thermophilic model organism *Parageobacillus thermoglucosidasius.* We engineered *P. thermoglucosidasius* to be strictly dependent for its growth on methanol assimilation via the core of the highly efficient ribulose monophosphate (RuMP) cycle, while co-assimilating ribose. Surprisingly, this did not require heterologous expression of RuMP enzymes. Instead, by laboratory evolution we awakened latent, native enzyme activities to form the core of the RuMP cycle. We obtained fast methylotrophic growth in which ∼17% of biomass was strictly obtained from methanol. This work lays the foundation for developing a versatile thermophilic bioproduction platform based on renewable methanol.

## Introduction

The use of microbial cell factories is a promising strategy to produce chemicals more sustainably. Several chemical products such as ethanol and lactate are already produced at a commercial scale by using this approach^1–5^. Sugar-based feedstocks are mostly used for the manufacturing of bio-based products, but there is an interest to move towards the use of alternative feedstocks to make bio-based manufacturing more sustainable and less dependent on agricultural land^6,7^. Alternative sustainable feedstocks should be generated from renewable resources and ideally not be dependent on agriculture, as this can potentially lead to competition with food production for production of bulk products, and a further reduction of the habitat for natural biodiversity.

Among alternative carbon feedstocks, one-carbon feedstocks such as formic acid, carbon monoxide and methanol are promising candidates, given that they can be produced from renewable resources. Among these feedstocks, methanol stands out because it is both water-miscible, electron rich, and its renewable production is already being scaled up. Methanol can be produced from biomass waste (bio-methanol) or from renewable electricity, CO_2_ and water (e-methanol)^6^. These characteristics make methanol suitable to sustainably produce the wide array of chemicals needed by society and appealing from a bioprocess and environmental perspective.

There are several pathways known to enable growth on methanol as a sole carbon and energy source, such as the serine cycle, the reductive glycine pathway, the xylulose monophosphate cycle or the ribulose monophosphate (RuMP) cycle. Among these pathways, the RuMP cycle is known to be one of the most efficient methanol assimilation pathways^6^. In all these pathways methanol needs to be oxidized first to formaldehyde via methanol dehydrogenase (Mdh). In the RuMP cycle, formaldehyde is then assimilated by condensation with ribulose 5-phosphate (Ru5P) to form hexulose 6-phosphate (H6P), in a reaction catalyzed by hexulose 6-phosphate synthase (Hps). H6P is isomerized to fructose 6-phosphate (F6P), via 6-phospho-3-hexuloisomerase (Phi). F6P can be further converted into biomass via several routes dependent on the exact variant of the RuMP cycle^6^. In the most ATP-efficient version of the RuMP cycle, the fructose bisphosphate aldolase/transaldolase (Fba/Tal) variant F6P is subsequently phosphorylated to fructose 1,6-bisphosphate (F1,6P), which is split via FBA into the correspondent trioses phosphate, glyceraldehyde 3-phosphate (G3P) and dihydroxyacetone phosphate (DHAP). A part of these triose phosphate molecules will be further metabolized following the glycolytic route, and a part of the G3P will be re-arranged via the pentose phosphate pathway (e.g. transketolase and transaldolase) to replenish the precursor Ru5P (Figure 1).

**Figure 1:**
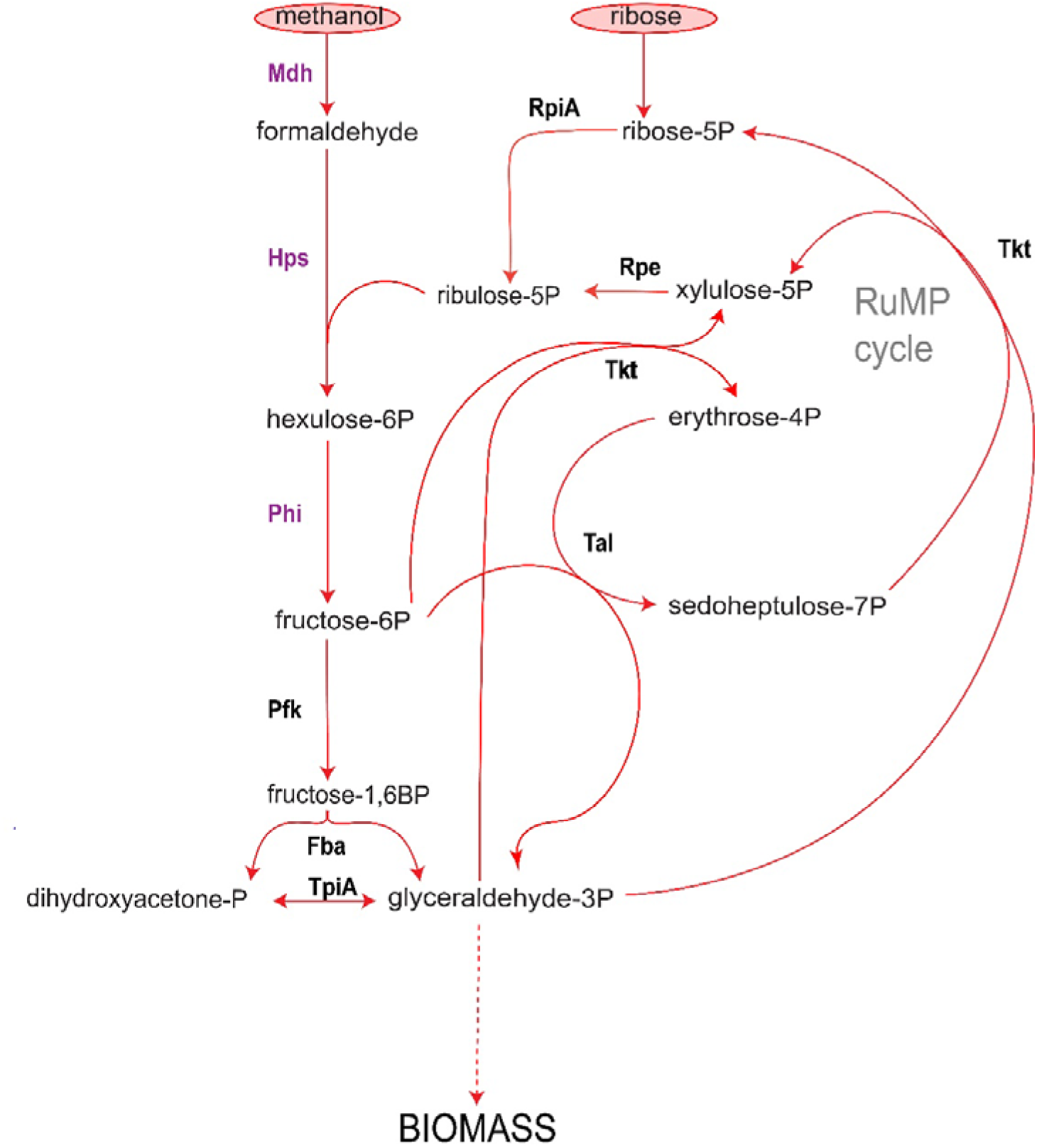
Metabolic pathway representing the most ATP-efficient variant of the RuMP cycle (Fba/Tal variant)^6^. The key enzymes for methylotrophy, Mdh, Hps and Phi are marked in purple. Mdh: methanol dehydrogenase, Hps: hexulose-6-phosphate synthase, Phi: 6-phospho 3-hexuloisomerase, RpiA: R5P isomerase, Rpe: Ru5P epimerase, Tkt: transketolase, Tal: transaldolase, Pfk: phosphofructokinase, Fba: F1,6P aldolase, TpiA: triose phosphate isomerase.

As methanol oxidation is a thermodynamically challenging reaction (Δ_r_G^’m^=30.5 kJ/mol)^8^, it represents a bottleneck for the implementation of methanol-based biotechnology^9,10^. However, it has been suggested that thermophilic organisms, which are known to have higher enzymatic rates, could be a solution to speed up the methanol oxidation bottleneck^11^. Moreover, the use of thermophilic organisms for biobased manufacturing can reduce the amount of energy needed to cool the bioreactors, as well as the risk of contamination by mesophilic organisms^12^. *Bacillus methanolicus* is the only well-known moderate thermophilic microorganism that can grow on methanol. It has an optimal growth temperature of 50°C and possesses the efficient RuMP cycle for growth on methanol as sole carbon and energy source (Figure 1)^13–15^. Even though *B. methanolicus* has been regarded and extensively studied as a promising thermophilic bacterial host for methanol-based manufacturing, it so far lacks effective molecular tools to modify its genome, which makes it a suboptimal microbial chassis for chemical production^16^.

It is a major bottleneck for the realization of thermophilic methanol-based bioproduction that there are limited thermophilic methylotrophic platform organisms with sufficient genetic tools. An alternative approach to obtain a suitable thermophilic microbial chassis for chemical production from methanol, is to engineer methylotrophy in a versatile, easier to handle thermophilic bacterium, with a broad range of genetic tools available. *Parageobacillus thermoglucosidasius* (formerly known as *Geobacillus thermoglucosidasius)* is a facultatively anaerobic thermophilic bacterium, which is emerging as a promising thermophilic platform organism. During the last years, an increasing number of genetic tools has been developed to effectively engineer its genome, which has harnessed its potential as a microbial chassis for the production of bulk and fine chemicals^17–20^. Overall, this makes *P. thermoglucosidasius* a promising thermophilic microbial chassis to engineer methylotrophy^17^

In the last decades, scientific efforts towards engineering synthetic methylotrophy in non-methylotrophic bacteria, yielded successful results in mesophilic model microorganisms. Full methylotrophy has been implemented in *Escherichia coli* and *Saccharomyces cerevisiae* ^21–27^, as well as partial methylotrophy in *Pseudomonas putida* and *Bacillus subtilis*^28,29^, but such life-style change has not been demonstrated yet for a thermophilic species. Engineering methylotrophy in a non-methylotrophic microorganism is a challenging process, which implies a holistic rewiring of central carbon and energy metabolism, as well as a readjustment of cellular homeostasis. Because of such complexity, typically efforts to engineer methylotrophy relied on a mixed approach, including rational engineering and adaptative laboratory evolution (ALE)^21–31^. These mixed approaches are often based on rational engineering of methanol-dependent strains as a starting point for ALE. Methanol-dependence can be achieved by one or more gene knockouts that make the cells unable to grow solely on a certain auxiliary carbon substrate unless they co-assimilate methanol. Such deletion strains are then normally complemented with heterologous enzymes that enable methanol and formaldehyde assimilation, such as Mdh and other enzymes. After methanol-dependency and heterologous enzymes are introduced, ALE is typically performed to enhance functional operation of methanol assimilation. Examples of rationally engineered methanol-dependent bacterial strains co-assimilating methanol and ribose (*rpe* deletion), xylose (*rpiA* deletion) or pyruvate (*tpiA or fba* deletions) can be found in literature ^22,23,25,28,31^. So far, in all cases that synthetic methylotrophy has been engineered in a non-methylotrophic organism, a heterologous Mdh was overexpressed to enable methanol oxidation in these strains ^21,23,26,29,32^. The most used Mdh’s in the field have been the ones from *B. methanolicus* (BmMdh), *Geobacillus stearothermophilus* (GsMdh), *Corynebacterium glutamicum* (CgMDH) and an engineered *Cupriavidus necator* N1 Mdh (CnMdh-CT4-1)^22,25,27,28,33,34^. In this work, we present the first instance of engineered partial methylotrophy via a native, promiscuous alcohol dehydrogenase.

In the present study, we used a mixed rational and evolutionary approach to engineer partial methylotrophy in a thermophilic bacterium for the first time, based on the methods and insights from previous efforts in mesophiles. However, unlike previous efforts in mesophiles, we could harness via laboratory evolution, the potential of the latent, native RuMP cycle enzymes without the expression of any heterologous enzymes. In these evolved strains of *P. thermoglucosidasius*, we demonstrated methanol-dependent growth with ribose co-assimilation. In addition, we discovered and benchmarked via *in vivo* assays a novel, thermophilic methanol dehydrogenase native to *P. thermoglucosidasius*. We believe this work is an important milestone in the path towards the realization of a thermophilic microbial chassis to produce green chemicals from methanol.

## Results

### *P. thermoglucosidasius* harbors both native formaldehyde oxidation and assimilation enzymes

To convert the non-methylotrophic *P. thermoglucosidasius* strain into a methylotroph, we started by exploring the native formaldehyde conversion enzymes. Formaldehyde is not only a central intermediate in the assimilation of methanol, but also a highly toxic metabolite as it can crosslink DNA and proteins at low concentrations^21^. Hence most organisms, including non-methylotrophic bacteria have developed different ways to convert formaldehyde into less toxic compounds.

A common way to detoxify formaldehyde found in microorganisms is its oxidation into formate and the subsequent oxidation of formate into CO_2_^35^. In *P. thermoglucosidasius* we identified a gene annotated as *adhP* (ALF09200.1), which encodes a protein with 37 % homology to the enzyme AdhA from *Bacillus subtilis*, a bacillithiol-dependent formaldehyde dehydrogenase with a detoxification role in this organism^28,35^. We hypothesized that if AdhP was a formaldehyde detoxification system in *P. thermoglucosidasius*, this protein would be upregulated upon addition of methanol as its (promiscuous) oxidation could lead to elevated formaldehyde levels in the cell. Analysis of the wild-type (WT) strain proteome revealed that AdhP was more than 18-fold overproduced when methanol was added (8 g/L; Figure 2), hinting that AdhP could, indeed be involved in formaldehyde oxidation and detoxification.

**Figure 2:**
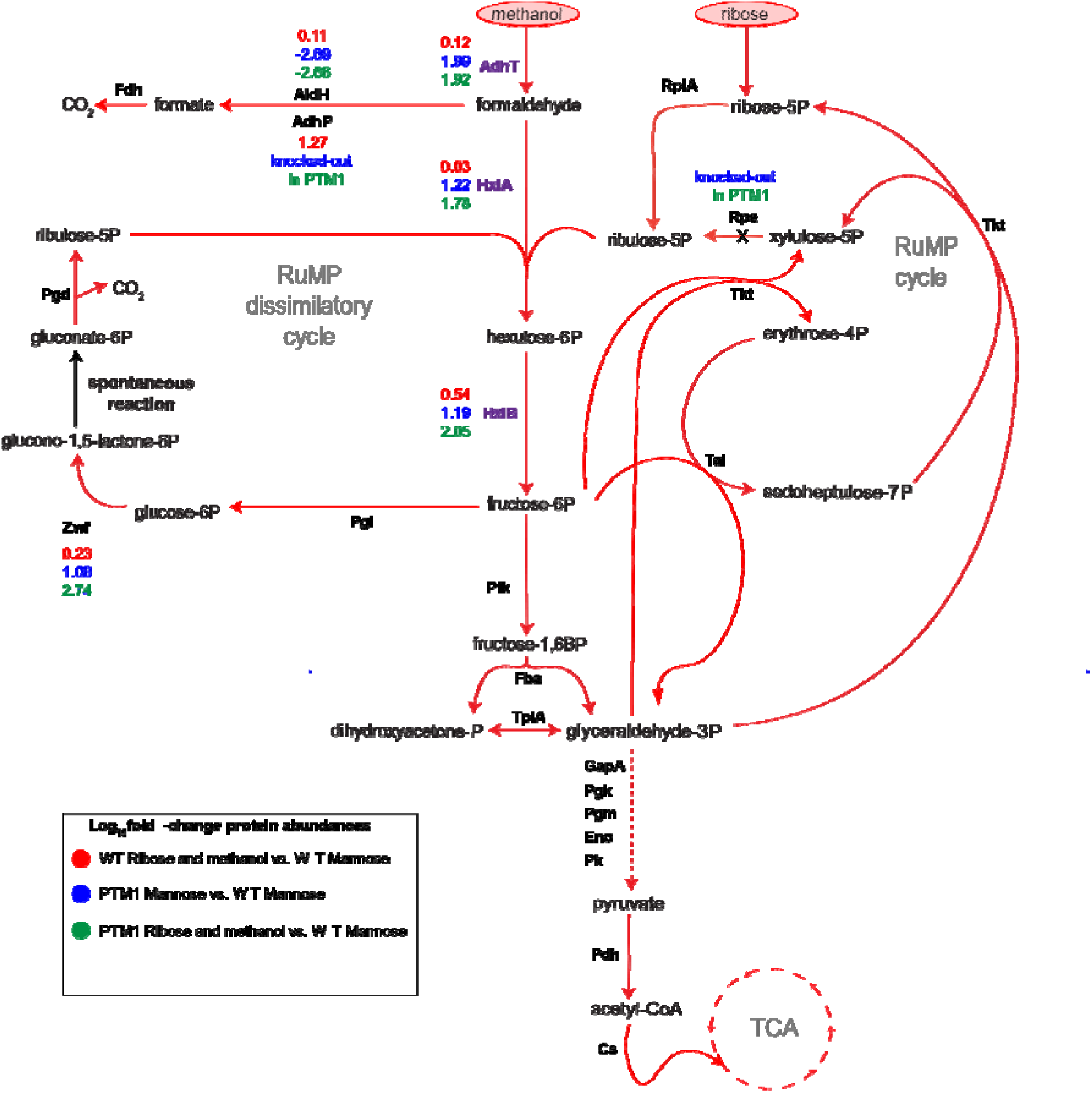
Metabolic pathway representing methanol and ribose co-assimilation via the RuMP cycle in the evolved partially methylotrophic strain PTM1 of *P. thermoglucosidasius*. Key native enzymes for methylotrophy, AdhT, Hps and Phi are in marked in purple. Coloured numbers indicate significant log_10_ fold-changes protein abundance levels between the WT strain and evolved PTM1 strain in different conditions. Proteins for which no numbers are indicated do not differ significantly in their abundance across conditions and strains. Volcano plots showing differential protein data for the full proteome can be found in Supplementary Figure S1 and full data are available in PRIDE (PXD058329).. AdhT: putative methanol dehydrogenase, Hps: hexulose-6-phosphate synthase encoded by *hxlA*, Phi: 6-phospho 3-hexuloisomerase encoded by *hxlB*, Zwf: G6P dehydrogenase, GapA: G3P dehydrogenase, RpiA: R5P isomerase, Rpe: R5P epimerase, Tkt: transketolase, Tal: transaldolase, Pfk: phosphofructokinase, Fba: fructose-1,6-bisphosphate aldolase, TpiA: triose phosphate isomerase, Pgk: phosphoglycerate kinase, Pgm: phosphoglycerate mutase, Eno: enolase, Pk: pyruvate kinase, Pdh: pyruvate dehydrogenase, Cs: citrate synthase, Pgi: phosphoglucose isomerase, Pgd: 6-phosphogluconate dehydrogenase, AdhP: putative formaldehyde dehydrogenase, Aldh: putative formaldehyde dehydrogenase.

As observed in several previous studies that established synthetic methylotrophy via the RuMP cycle, formaldehyde oxidation usually needs to be eliminated^21,23–29,32^. Hence, we deleted the gene encoding AdhP by homologous recombination and using a ThermoCas9 counterselection system^36^. We tested if there was an increased sensitivity to different concentrations of formaldehyde in the Δ*adhP* strain in comparison to the WT, however no differences were found (Table S1). We hypothesized that this dissonance between the results obtained from the proteome analysis and the similar formaldehyde sensitivity for the Δ*adhP* strain, could be explained by the existence of complementary formaldehyde detoxification systems in *P. thermoglucosidasius*.

In *B. subtilis*, two more formaldehyde detoxification systems are known. One of these systems involves the enzymes HxlA and HxlB, which are homologues of the RuMP enzymes Hps and Phi^35,37,38^. These enzymes can assimilate formaldehyde, by condensation with Ru5P and its further conversion into the central metabolite F6P. *P. thermoglucosidasius* may also have this capacity as it natively encodes HxlA and HxlB proteins (with 77.6 % and 76.7 % amino acid homology to the *B. methanolicus* proteins). We observed that, while HxlA was not upregulated in the *P. thermoglucosidasius* WT proteome during growth in the presence of methanol, HxlB was three-fold upregulated in the presence of methanol (Figure 2). This is surprising as both are encoded in the same operon and it is unclear why only second gene in the operon (*hxlB)* is higher expressed in the presence of methanol in the WT strain. We hypothesized that this assimilation route via HxlAB may also play a role in formaldehyde detoxication in *P. thermoglucosidasius*. Based on this knowledge, we aimed to exploit the native formaldehyde assimilation enzymes of *P. thermoglucosidasius* to harness methanol assimilation. As AdhP may convert formaldehyde into formate and CO_2_, which would divert carbon from biomass synthesis and bioproduction, we decided to use the *P. thermoglucosidasius* Δ*adhP* strain to further establish methylotrophy.

Awakening the latent RuMP cycle enzymes in *P. thermoglucosidasius* via the evolution of a methanol-dependent strain.

To realize methanol assimilation via the RuMP in *P. thermoglucosidasius,* we developed a strain that is strictly dependent on methanol assimilation during growth on ribose (Figure 3A). Hereto, we disrupted a part of the native ribose assimilation route, by deleting the gene *rpe*, which encodes for the enzyme ribulose-5-phosphate-3-epimerase. Rpe catalyzes the interconversion of ribulose-5-phosphate to xylulose-5-phosphate, and its activity is essential for growth on ribose as the sole carbon source. The Δ*adhP* Δ*rpe* strain was obtained with the ThermoCas9 counterselection tool, and we confirmed that this strain could not grow with ribose as a sole carbon source. We could then use this strain to develop a methanol-assimilating strain.

**Figure 3:**
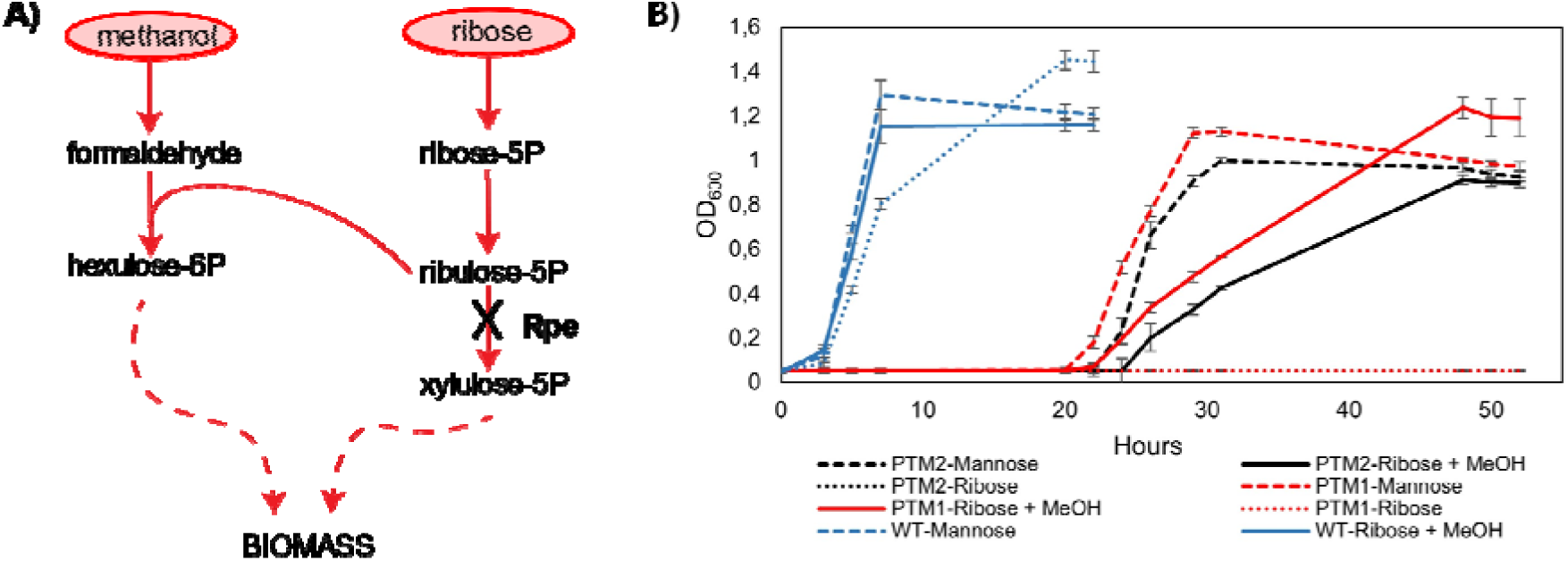
A) Methanol-dependency selection scheme dictated by the *rpe* gene deletion and used to evolve methylotrophy with ribose co-assimilation. B) Growth curves of the WT, the PTM1 and PTM2 of *P. thermoglucosidasius* cultivated in minimal media supplemented with different carbon sources. These results confirmed the methanol dependency of the evolved strains PTM1 and PTM2.

Methanol oxidation, the first step of methanol assimilation, is a crucial step to engineer methylotrophy. Typically, in synthetic methylotrophy a heterologous Mdh is introduced to enable methanol oxidation ^22,23,28,30,31^. *P. thermoglucosidasius* does not possess an annotated Mdh, however it possesses many alcohol dehydrogenases that could potentially take this role. Alcohol dehydrogenases are known to be promiscuous enzymes, which accept diverse types of alcohols as substrates. An example of this is the methanol dehydrogenase from *B. methanolicu*s MGA3, which has been demonstrated to oxidize propanol, butanol, and ethanol, even at higher rates than methanol itself ^39^. Thus, we hypothesized that via evolution of the Δ*adhP* Δ*rpe* methanol-dependent strain we could potentially exploit a promiscuous side activity of a native alcohol dehydrogenase from *P. thermoglucosidasius*, to take up the role of a methanol dehydrogenase.

Together with the natively present formaldehyde assimilation system encoded by the operon *hxlAB*, we hypothesized that we could evolve *P. thermoglucosidasius* Δ*adhP* Δ*rpe* to oxidize and assimilate methanol into biomass. To do this, we used the selective pressure dictated by the inability of the Δ*adhP* Δ*rpe* strain to grow on ribose as sole substrate. The methanol dependent *P. thermoglucosidasius* Δ*adhP* Δ*rpe* was cultivated in minimal TMM medium in the presence of ribose (4 g/L) with or without methanol (8 g/L). After three weeks, growth was observed in cultures containing methanol and ribose, but not in the ones containing only ribose. Two methanol-dependent strains were isolated from this short-term lab evolution culture and designated PTM1 and PTM2.

This result suggested that the native enzymatic machinery of the Δ*adhP* Δ*rpe* strain had evolved to co-assimilate ribose and methanol. To confirm the methanol dependency of the evolved PTM1 and PTM2 strains, we cultivated these strains together with the WT *P. thermoglucosidasius* in shake flasks and monitored their growth in the presence of either mannose, ribose, or a mixture of ribose and methanol as carbon sources (Figures 3B). After an extended lag-phase, both PTM1 and PTM2 could grow in the presence of ribose and methanol, while not on ribose alone. The cause of this lag phase is unknown to us, but is likely not related to the growth on methanol as also on mannose both strains show an extended lag phase. These results suggested that the evolved *P. thermoglucosidasius* strains assimilated methanol into biomass via native enzymes.

### Validating methanol assimilation via ^13^C labelling experiments

To validate methanol assimilation into biomass, we performed labelling experiments with ^13^C-labelled methanol. The evolved partially methylotrophic strains PTM1 and PTM2 were cultivated in minimal TMM medium with ^12^C-ribose (4 g/L) and ^13^C-labelled methanol (8 g/L). Cell pellets were analyzed to investigate the incorporation of the ^13^C-labelled methanol in the proteogenic amino acids serine and alanine. 3-phosphoglycerate (3PG) is the precursor of serine, while pyruvate is the precursor of alanine (Figure 4A). 3PG is a glycolytic intermediate and pyruvate the product of glycolysis, therefore these two metabolites are derived from the split of F6P into two triose phosphate molecules. We hypothesized that if methanol was co-assimilated with ribose via the RuMP enzymes, one of the carbon atoms of F6P should be labelled. Thus, 50% of the 3PG and pyruvate molecules should contain one labelled carbon atom, and so should their corresponding amino acid products, serine, and alanine (Figure 4A). The results of the analysis revealed that in the strain PTM1 ∼47 % of the serine and alanine molecules contained one ^13^C atom, matching well the expected values (Figure 4B). The labelling results for the strain PTM2 showed slightly lower labelling, being ∼39% and 38% for serine and alanine. The latter results on PTM2 suggested that this evolved strain possibly had leaky conversion of ribose into biomass independent from methanol assimilation. Overall, we concluded that these results validated methanol assimilation via the RuMP cycle enzymes in the partially methylotrophic evolved strains PTM1 and PTM2, and according to these results, we calculated that methanol accounts approximately 1/6^th^ (∼17%) of the biomass in the strain PTM1. This percentage was calculated taking into account that one carbon atom out of 6 in the main biomass precursor F6P comes from methanol assimilation, and the other 5 from ribose.

**Figure 4:**
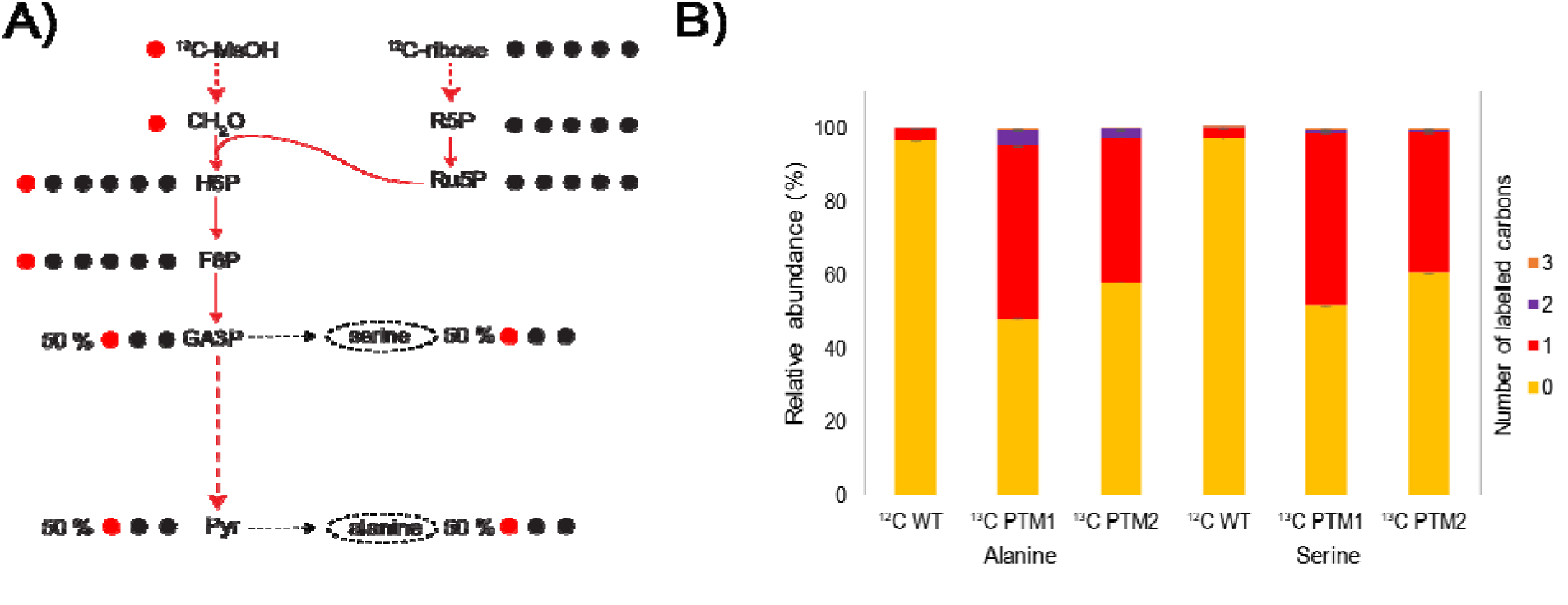
A) Metabolic scheme showing the expected labelling pattern of serine and alanine assuming co-assimilation of ^12^C-ribose and ^13^C-methanol B) Relative abundance of ^13^C-labelled carbons present in the proteogenic amino acids alanine and serine, in cells of the partially methylotrophic evolved strain PTM1 cultivated in the presence of a mixture of natural ^12^C ribose and ^13^C-labelled methanol. The WT strain cultivated on natural ^12^C ribose was used as control to account for naturally occurring ^13^C atoms. The data presented in this figure is based on biological duplicates.

### Whole genome sequencing of the evolved strains revealed crucial mutations for methanol assimilation

The genomes of the evolved partially methylotrophic *P. thermoglucosidasius* strains PTM1 and PTM2, as well as the parental strain Δ*adhP* Δ*rpe*, were sequenced and analyzed for mutations that occurred during the evolution towards partial methylotrophy. A list with all mutations found in the evolved strains can be found in the supplementary material (Table S2). We hypothesize that some of the overlapping mutations between these mutations can be related to the methylotrophic phenotype.

In relation to methanol oxidation, we observed that both strains had an insertion of 1447 bp in the upstream of the coding region of *adhT* gene, an uncharacterized alcohol dehydrogenase from *P. thermoglucosidasius.* We observed that this insertion corresponded to a mobile element (*tra905)* present in the genome of *P. thermoglucosidasius.* We hypothesized that AdhT is a promiscuous alcohol dehydrogenase with methanol dehydrogenase capabilities and this insertion upstream of the coding sequence increased transcription and/or translation of this enzyme to allow faster methanol conversion via AdhT.

Moreover, both evolved strains had a mutation (G->A) in codon 26 of the *hxlR* gene, resulting in an amino acid substitution (M26I). HxlR is a helix-turn-helix (HTH) transcriptional regulator controlling the expression of the operon *hxlAB,* which encodes central enzymes of the RuMP cycle. We hypothesized that the M26I mutation could be enhancing the expression of *hxlA* and *hxlB* and leading to an upregulation of these key enzymes for methanol assimilation.

Furthermore, the strains PTM1 and PTM2 have a large 23 KB deletion involving the whole *rbsDACBK* ribose operon and other genes. This seems a counterintuitive mutation as these strains were evolved while co-utilizing ribose. In the genome of *P. thermoglucosidasius* there is a second copy of the ribose transporter encoding the proteins *rbsABC*, but the genes *rbsD* and *rbsK,* which encode the proteins activating ribose to R5P, are not duplicated. Ribose activation is a necessary step for ribose and methanol co-assimilation, which indicates that this reaction can be possibly catalyzed by other enzymes. However, this remains an open question we could not answer in this study.

The *ptsG* gene encoding a glucose phosphotransferase system, was truncated in both PTM1 and PTM2 evolved strains, which could potentially block the uptake of glucose, but there is no obvious direct link between this and the partial methylotrophic growth phenotype.

Another interesting mutation that may have a plausible link to the RuMP operation was found only in PTM1. In this strain we found a SNP (1984003 T>A), which caused an amino acid substitution in the coding sequence of the *tkt* gene (T184A). *tkt* encodes the transketolase, an enzyme which catalyzes two of the reactions of the RuMP cycle (G3P + F6P = E4P + Xu5P; G3P + S7P = X5P + R5P; Figure 1). The Tyr184 residue is highly conserved among different species of microorganisms, and given its vicinity to the cofactor thiamine diphosphate binding site, this residue has been regarded as essential for the transketolase activity^40–42^. The activity of Tkt is essential for methanol and ribose co-assimilation via the RuMP, and therefore we believe this amino acid substitution did not result in a death variant. However, what is the actual effect of this mutation remains an open question could be studied in future studies via enzymatic assays

### Proteomics analysis revealed that key enzymes for the RuMP cycle were upregulated

To investigate the effect of some of the mutations observed in the *P. thermoglucosidasius* evolved strains, we aimed to analyze their proteome and compared it with the one of the *P. thermoglucosidasius* WT strain (Figure S1). Because the strain PTM2 showed lower methanol assimilation in the ^13^C-labelling, we decided to only continue analyzing the proteome of the PTM1 strain. The PTM1 and WT strains were cultivated in the presence of either mannose or a mixture of ribose and methanol as carbon sources.

In relation to the methanol oxidation and assimilation capabilities, we confirmed that the putative methanol dehydrogenase AdhT, as well as HxlA and HxlB were significantly upregulated in the PTM1 strain in comparison to the WT (Figure 2). AdhT was 98-fold (log_10_ fold change 1.99) more abundant in the PTM1 strain than in the WT,when both grown on mannose medium. This result confirmed our hypothesis that the mutations observed in the upstream region of the *adhT* gene, in PTM1, increased the expression of this alcohol dehydrogenase. The enzyme HxlA was more than ∼16-fold (log_10_ 1.22) overproduced in the PTM1 strain in comparison with the WT when grown on mannose, and even ∼60-fold (log_10_ 1.78) hen a mixture of ribose and methanol was used as carbon source compared to WTn grown on mannose. The enzyme HxlB was more than ∼15-fold (log_10_ 1.19) overproduced in the evolved PTM1 strain in comparison with the WT, when both grown on mannose. When PTM1 was grown on amixture of ribose and methanol HxlB was overproduced ∼112-fold (log_10_ 2.05) compared to the WT grown on mannose. These results indicated that the mutation observed in the HxlR transcriptional regulator in the evolved partially methylotrophic strains, increased the constitutive expression of the operon *hxlAB* in PTM1 without methanol present and further boosted the expression with methanol, thus enabling efficient assimilation of formaldehyde into fructose-6-phosphate. In conclusion, this strongly suggest that the native AdhT, HxlA and HxlB drive methanol assimilation in *P. thermoglucosidasius*.

In *B. subtilis*, it was shown by X-ray crystallography that HxlR undergoes a conformational change induced by the presence of formaldehyde, which promotes the formation of an intra-helical cysteine-lysine bond, and allows DNA binding to activate the expression of *hxlAB*^37,43^. The domains of the HxlR protein from *B. subtilis* and *P. thermoglucosidasius* responsible for this conformational change and activation of the expression of *hxlAB* are conserved. When comparing the corresponding amino acid residue 26 of HxlR, which was mutated in the evolved *P. thermoglucosidasius* strains PTM1 and PTM2, we have observed that this amino acid is situated in the formaldehyde recognition pocket, exactly in front of the cysteine-lysine bridge that activates HxlR. In light of this observation, we hypothesize that the M26I substitution could lead to a partial conformational change in the off state of HxlR, potentially causing the increased constitutive expression of HxlAB in the absence of the inducer formaldehyde observed in the evolved strain PTM1 (Figure 2). This hypothesis could explain why in the evolved strain PTM1, HxlA and HxlB are produced in higher amounts even when methanol is not present in the media.

In addition, we found that the enzyme glucose-6-phosphate dehydrogenase (Zwf), was more than 550-fold (log_10_ 2.74) overproduced in the evolved strain on ribose + methanol compared to the WT on mannose (Figure 2). This enzyme is part of the oxidative part of the RuMP cycle that diverts carbon from biosynthesis to produce NADPH and CO_2_, named the RuMP dissimilatory cycle. This dissimilatory cycle includes the reactions catalyzed by the enzymes G6P isomerase (Pgi), which isomerizes F6P into G6P, G6P dehydrogenase (Zwf), which oxidizes G6P into glucono-1,5-lactone-6P and regenerates NADPH, and 6-phosphogluconate dehydrogenase (Pgd), which dehydrogenase gluconate-6P into the precursor Ru5P and CO_2_ and also regenerates NAD(P)H(Figure 2). This result could indicate that the flux of carbon going through the RuMP dissimilatory pathway is considerable. This could cause that part of the carbon assimilated via formaldehyde condensation gets oxidized to CO_2_. Such dissimilatory route could support extra NAD(P)H regeneration or acts as a metabolic valve to avoid too high formaldehyde accumulation, as also observed in the native methylotroph *B. methanolicus*^44^. However, a too high dissimilatory flux may be wasteful, hence optimization of this flux may be helpful to obtain full methylotrophic growth. Also it should be noted that only Zwf seems to be upregulated and not the other two enzymes of the dissimilatory RuMP cycle (Pgi, Pgl), but it could be these are already present at high enough levels to carry enough flux for dissimilatory RuMP cycle activity.

Another interesting highly downregulated enzyme in the PTM1 proteome compared to the WT, was an enzyme potentially related to formaldehyde dissimilative detoxification. The enzyme annotated as aldehyde dehydrogenase (AldH), was ∼457-fold (log_10_ -2.66) downregulated in the evolved partially methylotrophic strain PTM1 relative to WT on mannose (Figure 2). In *Corynebacterium glutamicum*, a similar enzyme (CAF20817), which shares ∼70% amino acid identity with AldH from *P. thermoglucosidasius*, was confirmed to be part of a dissimilative formaldehyde detoxification pathway^35,45,46^. In *C. glutamicum* this pathway involved the oxidation of formaldehyde to formic acid by the acetaldehyde dehydrogenase and the dissimilation of formate to CO_2_. Thus, we hypothesize that the native AldH of *P. thermoglucosidasius* could be fulfilling a similar role, and the observed downregulation of AldH in the evolved PTM1 strain increases the pool of formaldehyde available for methylotrophic biosynthesis.

For the differential protein levels of AdhP and Zwf there is no direct clear mutations identified in the genome sequencing of the PTM1 strain, but indirect effects due to other mutations in regulatory proteins (e.g. PtsG) could possibly cause these drastic changes in protein abundance.

### *In vivo* assays confirmed that AdhT is a novel thermophilic methanol dehydrogenase

To validate that AdhT, the putative native methanol dehydrogenase from *P. thermoglucosidasius* indeed possessed methanol oxidation capabilities, we investigated the production of formaldehyde by this enzyme. We utilized two different *in vivo* approaches to detect the activity of AdhT, as methanol dehydrogenase activity is notoriously hard to measure *in vitro* due to the very low kinetic rates of this enzyme with methanol ^39^

In our first *in vivo* approach we confirmed the enzyme activity at thermophilic temperature (60°C) using plasmid-based overexpression of AdhT in *P. thermoglucosidasius* WT. In the second approach we further confirmed its activity and compared it to other commonly used Mdh’s at mesophilic temperature (37°C) in an *E. coli* formaldehyde biosensor strain^47^.

For the first approach, *P. thermoglucosidasius* strains overexpressing *adhT* from a plasmid under the transcriptional control of a medium strength promoter (*P. thermoglucosidasius* P1-*adhT*) and a strong promoter (*P. thermoglucosidasius* P2-*adhT*) were cultivated in the presence of methanol, and formaldehyde concentration was measured in the culture supernatants by a colorimetric method^48,49^ The formaldehyde concentration detected in *P. thermoglucosidasius* culture supernatants overexpressing the *adhT* gene under medium-strength P1 and strong P2 promoters, was respectively 15-fold and 19-fold higher. These results confirmed that the overexpression of the native AdhT enzyme from *P. thermoglucosidasius* enabled methanol oxidation, and hence AdhT can act as thermophilic methanol dehydrogenase. Formaldehyde was also detected in low concentrations in the supernatants of cultures of *P. thermoglucosidasius* harbouring an empty plasmid (Figure 5). This observed modest native methanol oxidation activity in *P. thermoglucosidasius* cultures, could be attributed to the presence of small amounts of AdhT expressed from the wild-type genome of *P. thermoglucosidasius* or the promiscuous activity of other alcohol dehydrogenases.

**Figure 5:**
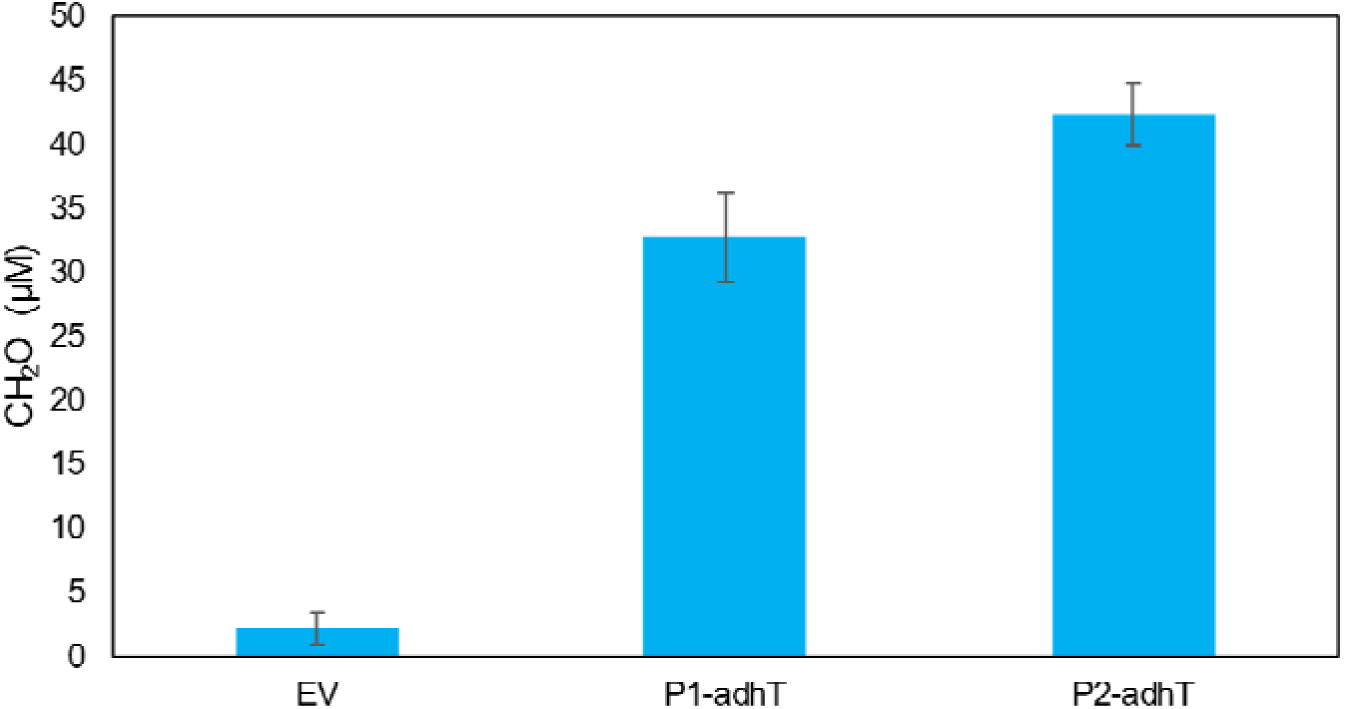
Average formaldehyde accumulation in culture supernatants of different *P. thermoglucosidasius* strains cultivated in the presence of methanol and overexpressing *adhT* under the control of the medium-strength promoter P1 (32.73 µM; *P. thermoglucosidasius* P1-*adhT*) and stronger promoter P2 (42.26 µM; *P. thermoglucosidasius* P2-*adhT*), or harboring an empty plasmid ( 2,14 µM; *P. thermoglucosidasius* EV).

The second *in vivo* approach to study the ability of AdhT to convert methanol into formaldehyde was performed in the *E. coli* HOB formaldehyde biosensor, published by Schann et al. (2024). This biosensor was engineered to be methionine and isoleucine auxotrophic. Growth of the biosensor can be rescued by the production of formaldehyde, which together with pyruvate can be converted to methionine and isoleucine via the combined activity of the 4-hydroxy-2-oxobutanoate (HOB) aldolase (HAL) and HOB transaminase (HAT) reactions^47^. In this experiment, either AdhT or other reference methanol dehydrogenase (CgMdh and BsMdh) were overexpressed in the *E. coli* formaldehyde biosensor and the resulting strains HOB–*Bsmdh,* HOB–*Cgmdh* and HOB–*adhT were* cultivated in varying concentrations of methanol. Growth of the biosensor implied that methanol was oxidized to formaldehyde, concluding that the overproduced Mdh possessed methanol dehydrogenase activity. When comparing AdhT performance to the other two tested Mdh’s, AdhT performed similar to the CgMdh, while BsMdh allowed better growth at lower methanol concentration (Figure 6). This likely indicates that AdhT has a relatively low affinity for methanol and required higher methanol concentrations to allow growth of the biosensor strain.

**Figure 6:**
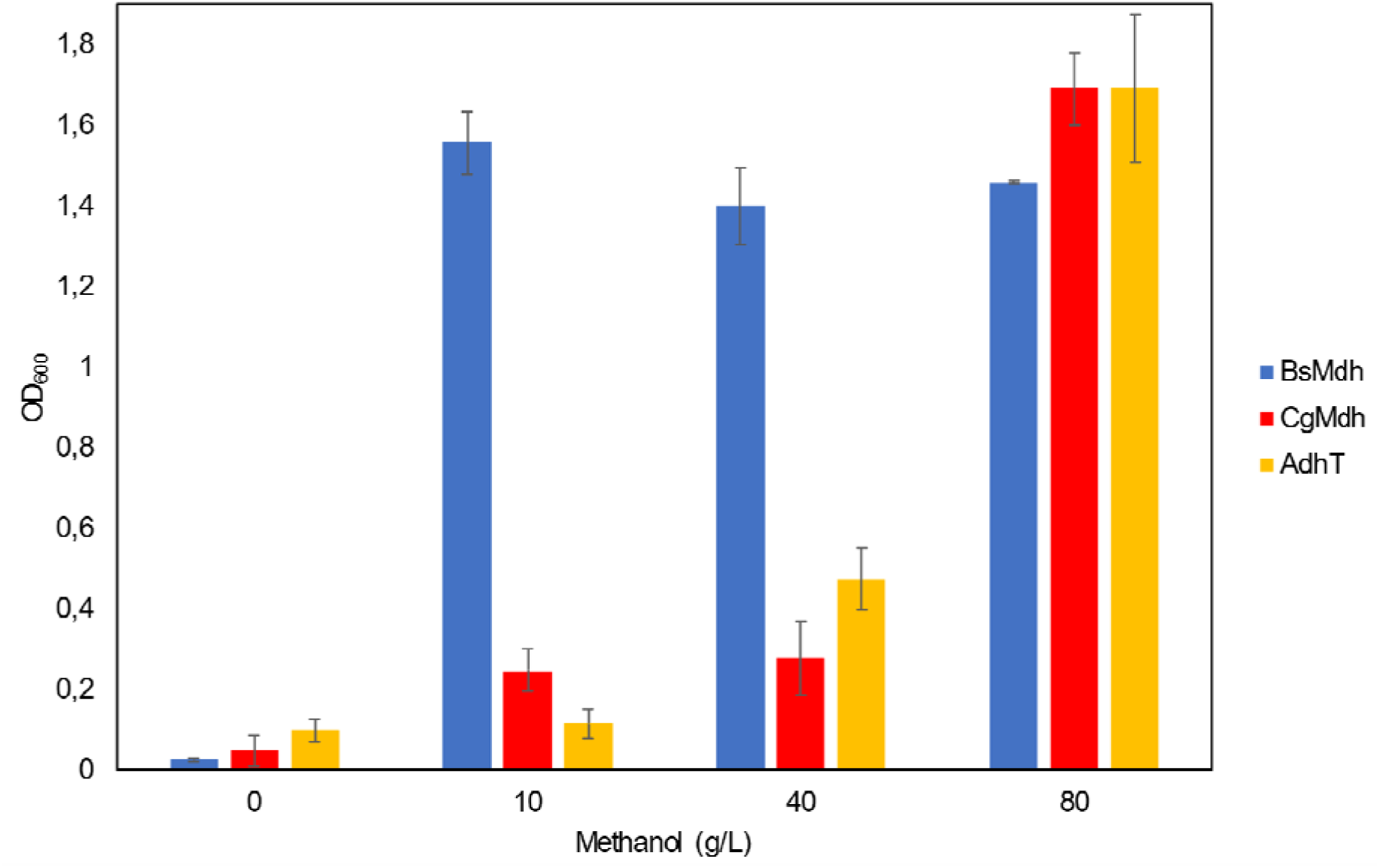
Average maximum OD_600_ of the *E. coli* formaldehyde HOB biosensor strain, overexpressing AdhT, BsMdh or CgMdh. The strains HOB –*Bsmdh,* HOB –*Cgmdh* and HOB –*adhT* were cultivated in M9 selective media supplemented with methanol in different concentrations.

Overall, the results from the mesophilic and thermophilic *in vivo* approaches demonstrated that the enzyme AdhT of *P. thermoglucosidasius* can be regarded as a newly discovered, thermophilic methanol dehydrogenase.

## Discussion

This work represents a milestone towards the realization of thermophilic methylotrophy in *P. thermoglucosidasius,* a promising thermophilic model organism for industrial use^17^. Our work presents the first example of engineering partial methylotrophy in a thermophilic organism, using a mixed rational and evolutionary approach. The present study also represents the first instance of engineering methylotrophy that was only reliant on activation of native latent RuMP cycle enzymes.

We identified key mutations the likely upregulate the protein abundances of native enzymes (AdhT, HxlA and HxlB,) that can catalyze reactions that are essential for methylotrophy via that RuMP cycle: Mdh, Hps and Hpi Other authors also demonstrated that during methylotrophic growth via the RuMP cycle, the Mdh, Hps and Phi enzymes were upregulated during evolution towards full methylotrophy and were key for methanol assimilation^23^. This was also demonstrated in the natural methylotroph *B. methanolicus*, in which adding extra copies of *hps* and *phi* (the homologues of *hxlAB*) increased the methanol tolerance and assimilation capacities of this bacterium^39,50^.

We furthermore confirmed the activity of AdhT as a novel thermophilic methanol dehydrogenase native to *P. thermoglucosidasius.*. We confirmed that this enzyme is a methanol dehydrogenase via two different *in vivo* assaysbased on the AdhT capacity to produce formaldehyde from methanol.

The ^13^C labelling experiment confirmed methanol assimilation in both isolated strains PTM1 and PTM2. However, the data of PTM2 showed that this strain had a lower percentage of once-labelled alanine and serine molecules than expected, and we hypothesized that this could be the consequence of some type of partial by-passing of the Rpe deletion,. Such a leaky metabolic phenotype has been reported previously for an *rpe* deficient *E. coli* strain^51^. Growth of this *rpe* knockout strain was observed 29 hours after inoculation on solely ribose and the authors hypothesized that *rpe* isozymes could be responsible for such an escape. However, after gene deletion of several putative *rpe* isozymes growth on solely ribose was not abolished and the cause of this leaky phenotype could not be identified. Also in this study we did not identify the cause of this phenotype in PTM2, however we note that for this strain we never observed sole growth on ribose. This together with the still large fraction of labelled amino acids (50% expected, ∼38% observed), indicates that the majority of the assimilation flux in PTM2 is still methanol-dependent.

Other studies have revealed that it is usual to find several formaldehyde detoxification systems some microorganisms. For instance, *B. subtilis* possesses three different formaldehyde detoxification systems. One based on the HxlAB enzymes (dissimilatory RuMP cycle), and two formaldehyde oxidation routes: the first one based on a BSH-dependent aldehyde dehydrogenase encoded by *adhA* (and another one of the recently discovered YycR aldehydedehydrogenase^52^. In *P. thermoglucosidasius,* we hypothesize that there are also three systems, HxlAB, AdhP and the uncharacterized aldehyde dehydrogenase encoded by *aldH*. We hypothesize that AldH detoxifies formaldehyde by oxidation into formate, which later gets dissimilated into CO_2_, in a similar way as AdhA in *B. subtilis.* We hypothesize that the drastic downregulation of AldH observed in the PTM1 strain in comparison with the WT, decreased the dissimilation of formaldehyde to CO_2_ and thus increased its availability for assimilation via HxlAB into biomass.

This study also sheds interesting light on potential evolutionary routes towards methylotrophy in nature. This work shows that only limited evolutionary steps are needed to evolve methylotrophy in a non-methylotroph. We observed that native enzymatic machinery, e.g. for alcohol oxidation, formaldehyde detoxification and pentose phosphate pathway can be activated with limited mutations towards functional methanol assimilation. It had been suggested previouslyfor the evolution of methylotrophy in some gram-negative bacteria, that methylotrophic growth may have evolved rather late in evolutionary history by (horizontally) combining different types of enzyme modules ^53^. For gram-positives, the only well-studied methylotroph is the thermophile *Bacillus methanolicus* ^54^. Interestingly, *P. thermoglucosidasius* is closely related to this species. This works hints that this feature in the *Bacilli* species may also have evolved possibly rather late in evolution. Most *Bacilli* are not methylotrophic, but our work shows that it could relatively easily evolve, especially as many *Bacilli* already possess Hps and Phi for formaldehyde detoxification, proving a good template for evolution of a full RuMP cycle. This may also be relevant for natural evolution of related species in niches where methylotrophy gives a fitness advantage.

We believe that the evolved strain PTM1 is a promising thermophilic platform starting point to establish full methylotrophy. To accomplish growth on methanol as a sole carbon source in this strain, first the *rpe* gene should be restored into the genome of the strain PTM1 to allow the full RuMP cycle to function. ALE is a common strategy to accomplish full methylotrophic growth after reconstitution of the full RuMP cycle in a partially methylotrophic strain ^23,25,31^. In this way, the strain is cultivated on a mixed carbon source including methanol and an auxiliary carbon source, and during a series of passages the auxiliary carbon source can be weaned out until growth on methanol as a sole carbon source is realized. In the case of the *P. thermoglucosidasius* PTM1 strain, after the *rpe* gene is reconstituted, it could be for example evolved with a mixture of methanol and ribose. The gene *aldH* could be a possible KO target to decrease wasteful losses of formaldehyde. Another potential target to study and possibly revert or alter for full methylotrophy would be the Tkt mutation in PTM1, as it is not clear if this mutation close to the active side is beneficial or detrimental for high flux through Tkt, which will be essential for full RumP cycle flux to support full methylotrophy.

Another strategy to generate a full methylotroph, which would also allow to better study evolutionary changes required for methylotrophy, could be to reverse engineer the mutations identified in this work. One could individually regenerate the mutations in a WT background and subsequently combine them to identify what is the causative set of mutations for partial methylotrophy. However, given the relatively time-consuming genetic engineering in *P. themorglucosidasius* this would require extensive experimental effort, and it may be more practical to use the PTM1 strain as starting point to generate a full methylotroph, as in fact was also done in recent successful efforts to generate synthetic methylotrophy in *E. coli* ^21,25^.

Overall this study paved the way towards a thermophilic methyltrophic platform organism. The strain generated in this work could be used for bioproduction from methanol, and further development of a full methylotrophic *P. thermoglucosidasius* for full methanol-based bioproduction.

## Materials and methods

### General cultivation techniques, media composition, bacterial strains, and plasmids

For the cultivation of *E. coli* strains, lysogeny broth (LB) medium was used containing 10 g/L tryptone (Bacto^TM^), 10 g/L NaCl, and 5 g/L yeast extract (Bacto^TM^). Solid LB medium additionally contained 15 g/L agar. *E. coli* strains were generally cultivated in 10 mL liquid medium in falcon tubes at 37°C and 200 rpm overnight or on solid medium at 37°C overnight. *E. coli* HOB biosensor cultures were supplemented with 0.25 mM of diaminopimelate acid (DAP).

For general cultivation of *P. thermoglucosidasius* strains, TGP medium was used containing 17 g/L tryptone (Bacto^TM^), 3 g/L peptone (Sigma Aldrich), 5 g/L NaCl, 2.5 g/L K_2_HPO_4_, 0.4% w/v glycerol, and 4 g/L sodium pyruvate. Solid TGP medium additionally contained 30 g/L agar. For the experimental investigation of methanol-dependent growth, formaldehyde concentration determination via NASH assay and the determination of growth curves of *P. thermoglucosidasius*, thermophilic minimal media (TMM) was used. TMM contained: 494 mg/L NaCl, 145 mg/L NaSO_4_, 24.7 mg/L KCl, 3.97 mg/L KBr, 184 mg/L MgCl_2_·6 H2O, 89.2 mg/L NaNO_3_, 8.37 g/L MOPS, 3.58 mg/L FeSO_4_, 716 mg/L tricine, 229 mg/L K_2_HPO_4_, 0.51 g/L NH_4_Cl, 73.5 mg/L CaCl_2_, 2 μg/L biotin, 5 μg/L thiamine-HCl, 2 μg/L folic acid, 0.1ug/L vitamin B12, 0.127 μg/L spermidine, 174 μg/L spermine tetrahydrochloride, 0.7 mg/L hemin, 1 mg/L FeCl_3_·6 H2O, 0.18 mg/L ZnSO_4_·7 H2O, 0.12 mg/L CuCl_2_·2 H2O, 0.12 mg/L, MnSO_4_· H2O, and 0.18 mg/L CoCl_2_·6 H2O, with a variable carbon source. Either D-ribose (4 g/L), glucose (5 g/L), mannose (10 g/L), methanol (8 g/L) or a mixture of sugars and methanol were added as carbon source. For the *in vivo* assays involving the *E. coli* HOB biosensor strain, M9 relaxing or selective media were used (Table S3).

When needed for cloning or plasmid propagation, chloramphenicol (15 µg/L), ampicillin (100 µg/L) or streptomycin (100 µg/L) was added to *E. coli* cultures, while either chloramphenicol (8 µg/L) or thiamphenicol (6.5 µg/L) was added to *P. thermoglucosidasius* cultures.

### Plasmid assembly and genome modification

Generally, primers were designed to either introduce BsaI sites and overhangs for Golden Gate assembly or to provide homology for Gibson assembly (Table S5). All the desired DNA fragments were amplified using Q5 polymerase (Q5 2x MasterMix, New England Biolabs), isolated from a 1% agarose gel (Zymogen gel DNA recovery kit) and assembled by either Gibson (50 °C, 1 hour) or Golden Gate ((37°C, 5 min; 16 °C, 5 min) x 30 ; 37°C, 20 min; 65°C, 20 min). 5 µL of assembly mixture were transformed by heat shock in Neb5a commercial chemically competent cells (New England Biolabs) and plasmid presence in bacterial cells was verified by colony PCR using OneTaq polymerase (OneTaq 2x MM, New England Biolabs). Plasmids were isolated (QIAprep Spin Miniprep Kit, Qiagen) and the sequence of all newly assembled plasmids were confirmed by Oxford Nanopore whole plasmid sequencing (Plasmidsaurus).

pZASS_*adhT* was constructed by Gibson assembly by combining two PCR amplified fragments with overlapping regions, one containing the *adhT* coding sequence, amplified from *P. thermoglucosidasius WT* genome as a template, and the other containing the pZASS backbone^47^. To obtain the plasmid pJS15A, a derivative of the Geobacilli:*E. coli* shuttle vector pJS28 (Table 1), we replaced by Golden Gate assembly the origin ColE1 from pJS28 by a lower copy number p15A ori ^57^. To construct the plasmids pJS15A-*adhP*KO and pJS15A-*rpe*KO, first *thermoCas9* under the expression of the *P. thermoglucosidasius* native cellobiose inducible promoter *pBgll*, and the sgRNA module including a scrambled spacer were amplified by PCR and assembled into the backbone pJS15A by Golden Gate. The resulting vector was later digested with SacI, and 1 KB homology arms (HA) flanking the upstream and downstream regions of either the *adhP* or *rpe* gene of *P. thermoglucosidasius* were included via Gibson assembly. The scrambled spacer was replaced by a targeting spacer via Gibson assembly.

**Table 1:** List of bacterial strains and plasmids used in this study, together with a brief description and the source.

| Bacterial strain or plasmid | Description | Source or reference |
| --- | --- | --- |
| *P. thermoglucosidasius* WT | Sporulation-deficient Type strain DSM 2542 with *sigF* gene deletion | <sup>55</sup> |
| *P. thermoglucosidasius* $\Delta adhP$ | *P. thermoglucosidasius* WT with *adhP* gene deletion | This work |
| *P. thermoglucosidasius* $\Delta adhP \Delta rpe$ | *P. thermoglucosidasius* $\Delta adhP$ with *rpe* gene deletion | This work |
| *P. thermoglucosidasius* PTM1 | Methanol dependent isolate evolved from *P. thermoglucosidasius* $\Delta adhP \Delta rpe$ | This work |
| *P. thermoglucosidasius* PTM2 | Methanol dependent isolate evolved from *P. thermoglucosidasius* $\Delta adhP \Delta rpe$ | This work |
| *P. thermoglucosidasius* P1-*adhT* | *P. thermoglucosidasius* WT harboring pJS15A-pRpsl8-*adhT* | This work |
| *P. thermoglucosidasius* P2-*adhT* | *P. thermoglucosidasius* WT harboring pJS15A-pLDH-*adhT* | This work |
| *P. thermoglucosidasius* EV | *P. thermoglucosidasius* WT harboring pJS15A empty vector | This work |
| *E. coli* NEB5a | *E. coli* cloning host.<br>*fhuA2Δ(argF-lacZ)* U169 *phoA*<br>*glnV44 Φ80Δ(lacZ)*M15 *gyrA96*<br>*recA1 relA1 endA1 thi-1 hsdR17* | New England Biolabs |
| *E. coli* HOB biosensor | Formaldehyde biosensor *E. coli*<br>$\Delta asd$, $\Delta frmRAB$, $\Delta rhmA$, *rhmA*. | <sup>47</sup> |
| HOB –*Bsmdh* | *E. coli* formaldehyde biosensor<br>harboring pZASS-*Bsmdh* | <sup>47</sup> |
| HOB –*Cgmdh* | *E. coli* formaldehyde biosensor<br>harboring pZASS-*Cgmdh* | <sup>47</sup> |
| HOB –*adhT* | *E. coli* formaldehyde biosensor<br>harboring pZASS-*adhT* | This work |
| pJS28 | Shuttle vector *Geobacillus-E. coli*. *repB*, *ColE1*, *ChlR*, *oriT*,<br>*truncated traJ* | <sup>55</sup> |
| pJS15A | Shuttle vector *Geobacillus-E. coli*. *repB*, *p15a*, *ChlR*, *oriT*,<br>*truncated traJ* | This work |
| pJS15A-*adhPKO* | pJS15A containing ThermoCas9,<br>a sgRNA targeting the *adhP*<br>gene of DSM 2542 | This work |
| pJS15A-*rpeKO* | pJS15A containing ThermoCas9,<br>a sgRNA targeting the *rpe* gene<br>of DSM 2542 | This work |
| pJS15A-pRpsl8-*adhT* | pJS15A containing *adhT* from *P. thermoglucosidasius*, under the<br>expression of pRpsl8 | This work |
| pJS15A-pLDH-*adhT* | pJS15A containing *adhT* from *P. thermoglucosidasius*, under the<br>expression of pLDH | This work |
| pZASS | *E. coli* overexpression plasmid<br>with p15A origin, streptomycin<br>resistance. | <sup>56</sup> |
| pZASS-*adhT* | pZASS containing *adhT* from *P. thermoglucosidasius*. | This work |

Confirmed plasmids were transformed into *P. thermoglucosidasius* by conjugation, following a slightly adapted protocol from Hon Wai Wan (2013)^58^. Notable differences were that a 50 mL culture of *P. thermoglucosidasius* was grown to mid-log phase (∼0.7 OD_600_) in a 250 mL baffled flask and then centrifuged at 4500xg for 10 minutes before the actual conjugation.

To delete the genes *adhP* and *rpe*, we induced homologous recombination of HA from pJS15A-*thermocas9*-*adhP*KO or pJS15A-*thermocas9*-*rpe*KO onto the genome of *P. thermoglucosidasius* WT strain and selected against wild-type DNA by using ThermoCas9. One confirmed transformant of *P. thermoglucosidasius* containing the editing plasmid was cultivated in 10 mL liquid TGP containing chloramphenicol (8 µg/mL) at 60°C overnight, then transferred (1mL) to fresh 10 mL TGP containing chloramphenicol (8 µg/mL) and cultivated at 68°C overnight. The resulting culture was then transferred into fresh TGP containing chloramphenicol (8 µg/mL) and 1% w/v cellobiose and cultivated at 60°C overnight to induce the *pBgl* promoter and enhance *thermocas9* transcription. Finally, the culture was plated on a TGP plate with 1% w/v cellobiose using 1 mL, 100 µL, 10 µL, or 1 µL of liquid culture and cultivated at 55°C. Gene deletions were confirmed by colony PCR.

Evolution of PTM partially methylotrophic strains of *P. thermoglucosidasius*.

To test if the *rpe* gene deletion indeed led to the inability to utilize ribose as a sole carbon source, we inoculated *P. thermoglucosidasius* Δ*adhP* Δ*rpe* in falcon tubes containing 10 mL of TMM supplemented with ribose in different concentrations (1, 2 and 4 g/L), and incubated them at 60°C and 180 rpm during at least two weeks. Growth was not observed in any of the cultures and therefore we concluded that this strain was not able to utilize ribose as a sole carbon source.

*P. thermoglucosidasius* Δ*adhP* Δ*rpe* was inoculated from glycerol stock in liquid TGP media and incubated overnight at 60°C and 180 rpm. Having previously removed the preculture media by centrifugation (5000 rpm, 3 minutes), 100 µL from the preculture were sub-cultured in a falcon tube containing 10 mL of TMM supplemented with ribose (4 g/L) and methanol (8 g/L). The cells were incubated at 60°C and 180 rpm for three weeks. After OD_600_ reached 0.6, methanol dependent growth was confirmed by subculturing the cells in 250 mL baffled flasks containing 25 mL of TMM supplemented with ribose (4 g/L) or with a mixture of ribose (4 g/L) and methanol (8 g/L). After 24h only the cultures containing methanol had grown, confirming that the evolved population was methanol dependent.

The evolved methanol-dependent *P. thermoglucosidasius* population was plated on TGP solid media from glycerol stocks and incubated at 60°C overnight. Up to ten colonies per strain were picked and screened for methanol-dependency in falcon tubes containing 10 mL of liquid TMM media containing ribose (4 g/L) or a mixture of ribose (4 g/L) and methanol (8 g/L). The isolates PTM1 and PTM2 which only grew when methanol was added, were selected for further analysis.

### Analysis of the genomic mutations occurring during evolution of partial methylotrophy

*P. thermoglucosidasius* genomic DNA was extracted from overnight cultures using the MasterPure™ Gram Positive DNA Purification Kit (Lucigen). The quality and concentration of the genomic DNA was verified by Qubit (ThermoFisher) and the genome was sequenced (Illumina, Novogene). The reads were mapped to a genome of reference and assembled in a contig, and the mutations differing between the evolved strains PTM1 and PTM2, and the parental strain were analyzed.

### Validation of methanol assimilation via ^13^C labelling experiments

To validate methanol assimilation into biomass, we performed steady-state labelling experiments with ^13^C labelled methanol. *P. thermoglucosidasius* WT, PTM1 and PTM2 strains were inoculated from glycerol stocks in liquid TGP media and incubated overnight at 60°C 180 rpm. 100 µL from each preculture were subcultured in 10 mL of TMM, containing natural labelled ribose (4 g/L) and supplemented with ^13^C-labelled methanol (8 g/L), having previously removed the preculture media by centrifugation (5000 rpm, 3 minutes). Cells were incubated at 60°C and 180 rpm, grown to late exponential phase (OD∼1), subcultured a second time in fresh media with ^13^C-labelled methanol and incubated overnight in the same conditions. One mL of culture broth was harvested (OD∼1), centrifuged (13000 rpm, 1 minute) and cell pellets were stored at -20 until further processing. Pellets were re-suspended in 200 µL of 6 M HCl and incubated at 110 °C for 16 hours to lyse the cells and hydrolyze the respective proteomes. Hydrolyzed samples were then filtered on a filter plate (AcroPrep Advance 96-well Filter, 0.2 µm membrane, product ID 8019) by positive pressure, and dried in a cold-trap speedvac. Dried samples were then derivatized by adding 35 µL of pyridine (Sigma-Aldrich, cat no. 270407) and 50 µL of MBTSFA + 1% (w/w) TBDMSClS (Sigma-Aldrich, cat no. 375934), mixed by pipetting and incubated at 60°C for 30 min. 75 µL of the derivatized samples were aliquoted to LC-MS vials with glass inserts for GC-MS analysis within 12h from derivatization on an Agilent 5977 GC-MS system with an Agilent DB-5ms capillary column (30m, inner diameter of 0.25 mm, film thickness of 0.25 µm, cat no. 122-5532). Samples were measured in full-scan mode, using a 1:100 split ratio, with the following gradient: start at 160°C, hold for 2 min, ramp to 310 °C at 7 °C/min, hold for 1 min. We considered for further analysis fragments listed in Long C.P. & Antoniewicz (2019) which included the whole C backbone of the relative amino acid (Ala 260, Ser 390)^59^. Raw chromatographic data was integrated using SmartPeak^60^. The relative data was corrected for the natural abundance of isotopes in the derivatization agents used for GCMS analysis^61^.

### Proteomic sample preparation

*P. thermoglucosidasius* PTM1 and WT strains were inoculated from glycerol stocks in liquid TGP media and incubated overnight at 60°C and 180 rpm. Having previously removed the preculture media by centrifugation (5000 rpm, 3 minutes), 100 µL from each preculture were subcultured in triplicated 250 mL baffled flasks containing 25 mL of TMM supplemented with either mannose (10 g/L) or a mixture of ribose (4 g/L) and methanol (8 g/L). Cells were incubated at 60°C and 180 rpm, grown to mid exponential phase (OD_600_=0.4-0.6), harvested by centrifugation (4700 rpm, 10 min), and stored at -20 until further processing. Samples were processed for mass spectrometry according to the SPEED protocol with slight modifications as described previously^62,63^. Shortly, cells were resuspended in 100 µL 100% TFA and transferred to clean plastic microtubes which were incubated in a thermocycler at 55 °C. A volume of 900 µL 2 M Tris solution (pH was not adjusted) was added and tubes were vortexed. The lysed samples were quantified by Bradford assay and stored at -80 °C. A mass of 20_μg of protein per sample was reduced (10_mM TCEP) and carbamidomethylated (55_mM CAA) for five_minutes at 95_°C. The proteins were digested by adding trypsin (proteomics grade, Roche) at a 1/50 enzyme/protein ratio (w/w) and incubated at 37°C overnight. Digests were acidified by the addition of 3% (v/v) formic acid (FA) and desalted using self-packed StageTips (five disks per micro-column, ø 1.5_mm, C18 material, 3_M Empore)^64^. The peptide eluates were dried to completeness and stored at −80_°C. Before the LC-MS/MS measurement, all samples were freshly resuspended in 12_µl 0.1% FA in HPLC grade water and 5 µl were subjected to LC-ESI-MS/MS.

### Proteomics data acquisition Exploris – microflow label-free DDA

Peptides were analyzed on a Vanquish Neo (microflow configuration) coupled to an Orbitrap Exploris 480 mass spectrometer (both Thermo Fisher Scientific). Around five micrograms of peptides were applied onto a commercially available Acclaim PepMap 100 C18 column (2 μm particle size, 1 mm ID × 150 mm, 100 Å pore size; Thermo Fisher Scientific) and separated using a stepped gradient with acetonitrile concentration ranging from 3% to 24% to 31% solvent B (0.1% FA, 3% DMSO in ACN) in solvent A (0.1% FA, 3% DMSO in HPLC grade water). A flow rate of 50 μl/min was applied. The mass spectrometer was operated in data-dependent acquisition (DDA) and positive ionization mode. MS1 full scans (360–1300 m/z) were acquired with a resolution of 60,000, a normalized automatic gain control target value of 100%, and a maximum injection time of 50 ms. Peptide precursor selection for fragmentation was carried out at a 1.2 seconds cycle time. Only precursors with charge states from two to six were selected, and dynamic exclusion of 30 s was enabled. Peptide fragmentation was performed using higher energy collision-induced dissociation and normalized collision energy of 28%. The precursor isolation window width of the quadrupole was set to 1.1 m/z. MS2 spectra were acquired with a resolution of 15,000, a fixed first mass of 100 m/z, a normalized automatic gain control target value of 100%, and a maximum injection time of 40 ms.

### LC-MS/MS data analysis

Peptide identification and quantification was performed using the software MaxQuant (version 1.6.3.4)^65^ with its built-in search engine Andromeda^66^. MS2 spectra were searched against the NCBI *P. thermoglucosidasius* protein database (GCF_001295365.1, downloaded May 2024), supplemented with common contaminants (built-in option in MaxQuant). Trypsin/P was specified as proteolytic enzyme. Carbamidomethylated cysteine was set as fixed modification. Oxidation of methionine and acetylation at the protein N-terminus was specified as variable modifications. Results were adjusted to 1% false discovery rate on peptide spectrum match (PSM) level and protein level employing a target-decoy approach using reversed protein sequences. Label-Free Quantification (LFQ)^67^ and iBAQ^68^ intensities were computed. The minimal peptide length was defined as seven amino acids and the “match-between-runs” functionality was disabled. The ProteinGroups.txt result file from MaxQuant was used as input for Omicsviewer 1.1.5^69^ to perform t-tests of differential protein abundances of *P. thermoglucosidasius* in the presence of mannose or ribose and methanol. Missing values were imputed by a protein-specific constant value, which was defined as the lowest detected protein-specific LFQ-value over all samples divided by two. Additionally, a maximal imputed LFQ value was defined as 15% quantile of the protein distribution from the complete dataset.

### Formaldehyde concentration estimation

*P. thermoglucosidasius* E, P1-*adhT* and P2-*adhT* strains were inoculated from glycerol stocks in liquid TGP media and incubated overnight at 60°C and 180 rpm. 100 µL from each preculture were subcultured in triplicated falcon tubes containing 10 mL of TMM supplemented with glucose (5 g/L), methanol (8 g/L) and thiamphenicol (6.5 mg/L) and incubated at 57°C and 180 rpm. One mL samples were taken at early stationary phase (OD_600_∼1) and formaldehyde concentration was estimated by using a colorimetric method^48^.

### *adhT* complementation *in vivo* experiment in *E. coli* HOB biosensor

*E.coli* HOB biosensor strains harboring the plasmids pZASS_*Bsmdh* and pZASS_*Cgmdh* were kindly donated by Schann et al. (2024). pZASS_*adhT* was cloned and transformed by heat shock in the *E. coli* HOB biosensor in house. Transformants of the HOB-*adhT* strain were confirmed via colony PCR as explained above. The strains HOB-*Bsmdh*, HOB-*Cgmdh* and HOB-*adhT* were streaked from glycerol stocks onto LB plates containing 0.25 mM DAP and streptomycin 100 mg/L, and grown overnight at 37°C. Individual colonies from each strain were inoculated in 10 ml M9 relaxing liquid medium (Table S 4) and incubated overnight at 37°C and 220 rpm. Cells were harvested and washed 3 times with M9 minimal medium at room temperature. The first washing step consisted of spinning down the cultures at 4700 rpm for 10 minutes, followed by resuspension in 900 μl. The second and third washing steps were performed at 1100 rpm for 1 minute. After washing was completed, experimental strains were diluted to a final OD_600_ of 0.01 in M9 selective media containing either 0, 10, 40 or 80 mM of methanol (Table S3). OD_600_ was measured using a portable ImplenTM OD_600_ DiluPhotometer. 150 μl of each culture was transferred to a transparent 96-well flat-bottomed plate with 50 μl mineral oil pipetted on top. For each condition and strain, three replicates were used. The growth of the different *E. coli* HOB strains was monitored for 70 hours in the BioTek synergy Neo2 multi-mode reader.

### Data availability

The mass spectrometric raw files as well as the MaxQuant output files have been deposited to the ProteomeXchange Consortium via the PRIDE partner repository and can be accessed using the identifier PXD058329^70^. gDNA raw reads corresponding to the PTM1, PTM2 strains and the parental Δ*adhP* Δ*rpe* strain have been deposited in the European Nucleotide Archive (accession number: PRJEB82962).

## Acknowledgements

We acknowledge Sebastian Wenk and his co-authors for supplying and providing advice on the use of the *E. coli* formaldehyde sensor strain. N.J.C. acknowledges support from his NWO Veni fellowship (VI.Veni.192.156). S.D. acknowledges support by The Novo Nordisk Foundation (NNF20CC0035580). The Exploris 480 mass spectrometer was funded in part by the German Research Foundation (INST 95/1435-1 FUGG). The authors would like to thank Chen Meng for use of the data visualization platform omicsViewer.

## References

1. Di Lorenzo, R. D., Serra, I., Porro, D., & Branduardi, P. (2022). State of the Art on the Microbial Production of Industrially Relevant Organic Acids. In Catalysts (Vol. 12, Issue 2). MDPI. 10.3390/catal12020234

2. Blombach, B., Grünberger, A., Centler, F., Wierckx, N., & Schmid, J. (2021). Exploiting unconventional prokaryotic hosts for industrial biotechnology. In Trends in Biotechnology. Elsevier Ltd. 10.1016/j.tibtech.2021.08.003

3. Liu, J., Han, X., Tao, F., & Xu, P. (n.d.). Reprogramming a high robust Geobacillus thermoglucosidasius 1 for efficient synthesis of polymer-grade lactic acid under 2 extremely high temperature (60° C). 10.1101/2023.04.14.536835

4. Zhou, J., Wu, K., & Rao, C. V. (2016). Evolutionary engineering of Geobacillus thermoglucosidasius for improved ethanol production; Evolutionary engineering of Geobacillus thermoglucosidasius for improved ethanol production. Biotechnol. Bioeng, 113, 2156–2167. 10.1002/bit.25983/abstract

5. Raita, M., Ibenegbu, C., Champreda, V., & Leak, D. J. (2016). Production of ethanol by thermophilic oligosaccharide utilising Geobacillus thermoglucosidasius TM242 using palm kernel cake as a renewable feedstock. Biomass and Bioenergy, 95, 45–54. 10.1016/j.biombioe.2016.08.015

6. Cotton, C. A., Claassens, N. J., Benito-Vaquerizo, S., & Bar-Even, A. (2020). Renewable methanol and formate as microbial feedstocks. In Current Opinion in Biotechnology (Vol. 62, pp. 168–180). Elsevier Ltd. 10.1016/j.copbio.2019.10.002

7. Liao, J. C., Mi, L., Pontrelli, S., & Luo, S. (2016). Fuelling the future: microbial engineering for the production of sustainable biofuels. Nature Reviews Microbiology, 14(5), 288–304. 10.1038/nrmicro.2016.32

8. Beber, M. E., Gollub, M. G., Mozaffari, D., Shebek, K. M., Flamholz, A. I., Milo, R., & Noor, E. (2022). eQuilibrator 3.0: a database solution for thermodynamic constant estimation. Nucleic Acids Research, 50(D1), D603–D609. 10.1093/nar/gkab1106

9. Heux, S., Brautaset, T., Vorholt, J. A., Wendisch, V. F., & Portais, J. C. (2018). Synthetic Methylotrophy: Past, Present, and Future. In Methane Biocatalysis: Paving the Way to Sustainability (pp. 133–151). Springer International Publishing. 10.1007/978-3-319-74866-5_9

10. Krüsemann, J. L., Rainaldi, V., Cotton, C. A., Claassens, N. J., & Lindner, S. N. (2023). The cofactor challenge in synthetic methylotrophy: bioengineering and industrial applications. Current Opinion in Biotechnology, 82, 102953. 10.1016/j.copbio.2023.102953

11. Whitaker, W. B., Sandoval, N. R., Bennett, R. K., Fast, A. G., & Papoutsakis, E. T. (2015). Synthetic methylotrophy: engineering the production of biofuels and chemicals based on the biology of aerobic methanol utilization. Current Opinion in Biotechnology, 33, 165–175. 10.1016/j.copbio.2015.01.007

12. Komives, C. F., Cheung, L. Y.-Y., Pluschkell, S. B., & Flickinger, M. C. (2005). Growth of Bacillus methanolicus in seawater-based media. Journal of Industrial Microbiology & Biotechnology, 32(2), 61–66. 10.1007/s10295-004-0195-9

13. Müller, J. E. N., Heggeset, T. M. B., Wendisch, V. F., Vorholt, J. A., & Brautaset, T. (2015). Methylotrophy in the thermophilic Bacillus methanolicus, basic insights and application for commodity production from methanol. In Applied Microbiology and Biotechnology (Vol. 99, Issue 2, pp. 535–551). Springer Verlag. 10.1007/s00253-014-6224-3

14. Müller, J. E. N., Meyer, F., Litsanov, B., Kiefer, P., & Vorholt, J. A. (2015). Core pathways operating during methylotrophy of Bacillus methanolicusMGA3 and induction of a bacillithiol-dependent detoxification pathway upon formaldehyde stress. Molecular Microbiology, 98(6), 1089–1100. 10.1111/mmi.13200

15. Brautaset, T., Jakobsen, Ø. M., Josefsen, K. D., Flickinger, M. C., & Ellingsen, T. E. (2007). Bacillus methanolicus: A candidate for industrial production of amino acids from methanol at 50°C. In Applied Microbiology and Biotechnology (Vol. 74, Issue 1, pp. 22–34). 10.1007/s00253-006-0757-z

16. Yang, X., Zheng, Z., & Wang, Y. (2024). Bacillus methanolicus: an emerging chassis for low-carbon biomanufacturing. Trends in Biotechnology. 10.1016/j.tibtech.2024.06.013

17. Paredes-Barrada, M., Kopsiaftis, P., Claassens, N. J., & van Kranenburg, R. (2024). Parageobacillus thermoglucosidasius as an emerging thermophilic cell factory. Metabolic Engineering, 83, 39–51. 10.1016/j.ymben.2024.03.001

18. Millgaard, M., Bidart, G. N., Pogrebnyakov, I., Nielsen, A. T., & Welner, D. H. (2023). An improved integrative GFP-based vector for genetic engineering of Parageobacillus thermoglucosidasius facilitates the identification of a key sporulation regulator. AMB Express, 13(1), 44. 10.1186/s13568-023-01544-9

19. Yang, Z., Li, Z., Li, B., Bu, R., Tan, G.-Y., Wang, Z., Yan, H., Xin, Z., Zhang, G., Li, M., Xiang, H., Zhang, L., & Wang, W. (2023). A thermostable type I-B CRISPR-Cas system for orthogonal and multiplexed genetic engineering. Nature Communications, 14(1), 6193. 10.1038/s41467-023-41973-5

20. Simon, M., & Msci, H. L. (n.d.). The Expansion of Available Bioparts and the Development of CRISPR/Cas9 for Genetic Engineering of Parageobacillus thermoglucosidasius.

21. Chen, F. Y. H., Jung, H. W., Tsuei, C. Y., & Liao, J. C. (2020). Converting Escherichia coli to a Synthetic Methylotroph Growing Solely on Methanol. Cell, 182(4), 933–946.e14. 10.1016/j.cell.2020.07.010

22. Kim, S., Lindner, S. N., Aslan, S., Yishai, O., Wenk, S., Schann, K., & Bar-Even, A. (2020). Growth of E. coli on formate and methanol via the reductive glycine pathway. Nature Chemical Biology, 16(5), 538–545. 10.1038/s41589-020-0473-5

23. Reiter, M. A., Bradley, T., Büchel, L. A., Keller, P., Hegedis, E., Gassler, T., & Vorholt, J. A. (2024). A synthetic methylotrophic Escherichia coli as a chassis for bioproduction from methanol. Nature Catalysis, 7(5), 560–573. 10.1038/s41929-024-01137-0

24. Nieh, L.-Y., Chen, F. Y.-H., Jung, H.-W., Su, K.-Y., Tsuei, C.-Y., Lin, C.-T., Lee, Y.-Q., & Liao, J. C. (2024). Evolutionary engineering of methylotrophic E. coli enables fast growth on methanol. Nature Communications, 15(1), 8840. 10.1038/s41467-024-53206-4

25. Keller, P., Reiter, M. A., Kiefer, P., Gassler, T., Hemmerle, L., Christen, P., Noor, E., & Vorholt, J. A. (2022). Generation of an Escherichia coli strain growing on methanol via the ribulose monophosphate cycle. Nature Communications, 13(1), 5243. 10.1038/s41467-022-32744-9

26. Whitaker, W. B., Jones, J. A., Bennett, K., Gonzalez, J., Vernacchio, V. R., Collins, S. M., Palmer, M. A., Schmidt, S., Antoniewicz, M. R., Koffas, M. A., & Papoutsakis, E. T. (n.d.). Engineering the Biological Conversion of Methanol to Specialty Chemicals in Escherichia coli.

27. Zhan, C., Li, X., Lan, G., Baidoo, E. E. K., Yang, Y., Liu, Y., Sun, Y., Wang, S., Wang, Y., Wang, G., Nielsen, J., Keasling, J. D., Chen, Y., & Bai, Z. (2023). Reprogramming methanol utilization pathways to convert Saccharomyces cerevisiae to a synthetic methylotroph. Nature Catalysis, 6(5), 435–450. 10.1038/s41929-023-00957-w

28. Gao, B., Zhao, N., Deng, J., Gu, Y., Jia, S., Hou, Y., Lv, X., & Liu, L. (2022). Constructing a methanol-dependent Bacillus subtilis by engineering the methanol metabolism. Journal of Biotechnology, 343, 128–137. 10.1016/j.jbiotec.2021.12.005

29. Bruinsma, L., Wenk, S., Claassens, N. J., & Martins dos Santos, V. A. P. (2023). Paving the way for synthetic C1 - Metabolism in Pseudomonas putida through the reductive glycine pathway. Metabolic Engineering, 76, 215–224. 10.1016/j.ymben.2023.02.004

30. Meyer, F., Keller, P., Hartl, J., Gröninger, O. G., Kiefer, P., & Vorholt, J. A. (2018). Methanol-essential growth of Escherichia coli. Nature Communications, 9(1). 10.1038/s41467-018-03937-y

31. Keller, P., Noor, E., Meyer, F., Reiter, M. A., Anastassov, S., Kiefer, P., & Vorholt, J. A. (2020). Methanol-dependent Escherichia coli strains with a complete ribulose monophosphate cycle. Nature Communications, 11(1). 10.1038/s41467-020-19235-5

32. Müller, J. E. N., Meyer, F., Litsanov, B., Kiefer, P., Potthoff, E., Heux, S., Quax, W. J., Wendisch, V. F., Brautaset, T., Portais, J. C., & Vorholt, J. A. (2015). Engineering Escherichia coli for methanol conversion. Metabolic Engineering, 28, 190–201. 10.1016/j.ymben.2014.12.008

33. Wu, T.-Y., Chen, C.-T., Liu, J. T.-J., Bogorad, I. W., Damoiseaux, R., & Liao, J. C. (2016). Characterization and evolution of an activator-independent methanol dehydrogenase from Cupriavidus necator N-1. Applied Microbiology and Biotechnology, 100(11), 4969–4983. 10.1007/s00253-016-7320-3

34. Lee, J. Y., Park, S. H., Oh, S. H., Lee, J. J., Kwon, K. K., Kim, S. J., Choi, M., Rha, E., Lee, H., Lee, D. H., Sung, B. H., Yeom, S. J., & Lee, S. G. (2020). Discovery and Biochemical Characterization of a Methanol Dehydrogenase From Lysinibacillus xylanilyticus. Frontiers in Bioengineering and Biotechnology, 8. 10.3389/fbioe.2020.00067

35. Klein, V. J., Irla, M., Gil López, M., Brautaset, T., & Fernandes Brito, L. (2022). Unravelling Formaldehyde Metabolism in Bacteria: Road towards Synthetic Methylotrophy. Microorganisms, 10(2), 220. 10.3390/microorganisms10020220

36. Mougiakos, I., Mohanraju, P., Bosma, E. F., Vrouwe, V., Finger Bou, M., Naduthodi, M. I. S., Gussak, A., Brinkman, R. B. L., Van Kranenburg, R., & Van Der Oost, J. (2017). Characterizing a thermostable Cas9 for bacterial genome editing and silencing. Nature Communications, 8(1). 10.1038/s41467-017-01591-4

37. Yurimoto, H., Hirai, R., Matsuno, N., Yasueda, H., Kato, N., & Sakai, Y. (2005). HxlR, a member of the DUF24 protein family, is a DNA-binding protein that acts as a positive regulator of the formaldehyde-inducible hxlAB operon in Bacillus subtilis. Molecular Microbiology, 57(2), 511–519. 10.1111/j.1365-2958.2005.04702.x

38. Yasueda, H., Kawahara, Y., & Sugimoto, S. (1999). Bacillus subtilis yckG and yckF Encode Two Key Enzymes of the Ribulose Monophosphate Pathway Used by Methylotrophs, and yckH Is Required for Their Expression. Journal of Bacteriology, 181(23), 7154–7160. 10.1128/JB.181.23.7154-7160.1999

39. Krog, A., Heggeset, T. M. B., Müller, J. E. N., Kupper, C. E., Schneider, O., Vorholt, J. A., Ellingsen, T. E., & Brautaset, T. (2013). Methylotrophic Bacillus methanolicus Encodes Two Chromosomal and One Plasmid Born NAD+ Dependent Methanol Dehydrogenase Paralogs with Different Catalytic and Biochemical Properties. PLoS ONE, 8(3). 10.1371/journal.pone.0059188

40. Kovina, M., Viryasov, M., Baratova, L., & Kochetov, G. (1996). Localization of reactive tyrosine residues of baker’s yeast transketolase. FEBS Letters, 392(3), 293–294. 10.1016/0014-5793(96)00836-8

41. Kovina, M. V, & Kochetov, G. A. (1998). Cooperativity and flexibility of active sites in homodimeric transketolase. FEBS Letters, 440(1–2), 81–84. 10.1016/S0014-5793(98)01423-9

42. Ga, K., & Ia, S. (2005). Binding of the Coenzyme and Formation of the Transketolase Active Center. IUBMB Life (International Union of Biochemistry and Molecular Biology: Life), 57(7), 491–497. 10.1080/15216540500167203

43. Zhu, R., Zhang, G., Jing, M., Han, Y., Li, J., Zhao, J., Li, Y., & Chen, P. R. (2021). Genetically encoded formaldehyde sensors inspired by a protein intra-helical crosslinking reaction. Nature Communications, 12(1), 581. 10.1038/s41467-020-20754-4

44. Delépine, B., López, M. G., Carnicer, M., Vicente, C. M., Wendisch, V. F., & Heux, S. (2020). Charting the Metabolic Landscape of the Facultative Methylotroph Bacillus methanolicus. MSystems, 5(5). 10.1128/mSystems.00745-20

45. Witthoff, S., Mühlroth, A., Marienhagen, J., & Bott, M. (2013). C 1 Metabolism in Corynebacterium glutamicum: an Endogenous Pathway for Oxidation of Methanol to Carbon Dioxide. Applied and Environmental Microbiology, 79(22), 6974–6983. 10.1128/AEM.02705-13

46. Lessmeier, L., Hoefener, M., & Wendisch, V. F. (2013). Formaldehyde degradation in Corynebacterium glutamicum involves acetaldehyde dehydrogenase and mycothiol-dependent formaldehyde dehydrogenase. Microbiology, 159(Pt_12), 2651–2662. 10.1099/mic.0.072413-0

47. Schann, K., Bakker, J., Boinot, M., Kuschel, P., He, H., Nattermann, M., Paczia, N., Erb, T., Bar-Even, A., & Wenk, S. (2024). Design, construction and optimization of formaldehyde growth biosensors with broad application in biotechnology. Microbial Biotechnology, 17(7). 10.1111/1751-7915.14527

48. Nash, T. (1953). The colorimetric estimation of formaldehyde by means of the Hantzsch reaction. Biochemical Journal, 55(3), 416–421. 10.1042/bj0550416

49. Reeve, B., Martinez-Klimova, E., De Jonghe, J., Leak, D. J., & Ellis, T. (2016). The Geobacillus Plasmid Set: A Modular Toolkit for Thermophile Engineering. ACS Synthetic Biology, 5(12), 1342– 1347. 10.1021/acssynbio.5b00298

50. Jakobsen, Ø. M., Benichou, A., Flickinger, M. C., Valla, S., Ellingsen, T. E., & Brautaset, T. (2006). Upregulated transcription of plasmid and chromosomal ribulose monophosphate pathway genes is critical for methanol assimilation rate and methanol tolerance in the methylotrophic bacterium Bacillus methanolicus. Journal of Bacteriology, 188(8), 3063–3072. 10.1128/JB.188.8.3063-3072.2006

51. Chen, C.-T., Chen, F. Y.-H., Bogorad, I. W., Wu, T.-Y., Zhang, R., Lee, A. S., & Liao, J. C. (2018). Synthetic methanol auxotrophy of Escherichia coli for methanol-dependent growth and production. Metabolic Engineering, 49, 257–266. 10.1016/j.ymben.2018.08.010

52. Klein, V. J., Troøyen, S. H., Fernandes Brito, L., Courtade, G., Brautaset, T., & Irla, M. (2024). Identification and characterization of a novel formaldehyde dehydrogenase in Bacillus subtilis. Applied and Environmental Microbiology, 90(11). 10.1128/aem.02181-23

53. Chistoserdova, L., Lapidus, A., Han, C., Goodwin, L., Saunders, L., Brettin, T., Tapia, R., Gilna, P., Lucas, S., Richardson, P. M., & Lidstrom, M. E. (2007). Genome of Methylobacillus flagellatus, Molecular Basis for Obligate Methylotrophy, and Polyphyletic Origin of Methylotrophy. Journal of Bacteriology, 189(11), 4020–4027. 10.1128/JB.00045-07

54. Brautaset, T., Jakobsen, Ø. M., Josefsen, K. D., Flickinger, M. C., & Ellingsen, T. E. (2007). Bacillus methanolicus: a candidate for industrial production of amino acids from methanol at 50°C. Applied Microbiology and Biotechnology, 74(1), 22–34. 10.1007/s00253-006-0757-z

55. Marinus Petrus Machielsen, Mariska Van Hartskamp, & Richard Van Kranenburg. (2015). Genetically modified (r)-lactic acid producing thermophilic bacteria (Patent WO2016012296A1).

56. Wenk, S., Yishai, O., Lindner, S. N., & Bar-Even, A. (2018). An Engineering Approach for Rewiring Microbial Metabolism (pp. 329–367). 10.1016/bs.mie.2018.04.026

57. Richard Van KranenburgMariska Van HartskampEelco Anthonius Johannes HeintzEsther Johanna Geertruda Van MullekomJurgen Snelders. (2013). Genetic modification of homolactic thermophilic bacilli.

58. Hon Wai Wan. (2013). Transformation of the thermophilic bacterium, Geobacillus debilis, by conjugation with the mesophilic bacterium, Escherichia coli.

59. Long, C. P., & Antoniewicz, M. R. (2019). High-resolution 13C metabolic flux analysis. Nature Protocols, 14(10), 2856–2877. 10.1038/s41596-019-0204-0

60. Kutuzova, S., Colaianni, P., Röst, H., Sachsenberg, T., Alka, O., Kohlbacher, O., Burla, B., Torta, F., Schrübbers, L., Kristensen, M., Nielsen, L., Herrgård, M. J., & McCloskey, D. (2020). SmartPeak Automates Targeted and Quantitative Metabolomics Data Processing. Analytical Chemistry, 92(24), 15968–15974. 10.1021/acs.analchem.0c03421

61. Wahl, S. A., Dauner, M., & Wiechert, W. (2004). New tools for mass isotopomer data evaluation in 13 C flux analysis: Mass isotope correction, data consistency checking, and precursor relationships. Biotechnology and Bioengineering, 85(3), 259–268. 10.1002/bit.10909

62. Abele, M., Doll, E., Bayer, F. P., Meng, C., Lomp, N., Neuhaus, K., Scherer, S., Kuster, B., & Ludwig, C. (2023). Unified Workflow for the Rapid and In-Depth Characterization of Bacterial Proteomes. Molecular & Cellular Proteomics, 22(8), 100612. 10.1016/j.mcpro.2023.100612

63. Doellinger, J., Schneider, A., Hoeller, M., & Lasch, P. (2020). Sample Preparation by Easy Extraction and Digestion (SPEED) - A Universal, Rapid, and Detergent-free Protocol for Proteomics Based on Acid Extraction. Molecular & Cellular Proteomics, 19(1), 209–222. 10.1074/mcp.TIR119.001616

64. Rappsilber, J., Mann, M., & Ishihama, Y. (2007). Protocol for micro-purification, enrichment, pre-fractionation and storage of peptides for proteomics using StageTips. Nature Protocols, 2(8), 1896– 1906. 10.1038/nprot.2007.261

65. Tyanova, S., Temu, T., & Cox, J. (2016). The MaxQuant computational platform for mass spectrometry-based shotgun proteomics. Nature Protocols, 11(12), 2301–2319. 10.1038/nprot.2016.136

66. Cox, J., Neuhauser, N., Michalski, A., Scheltema, R. A., Olsen, J. V., & Mann, M. (2011). Andromeda: A Peptide Search Engine Integrated into the MaxQuant Environment. Journal of Proteome Research, 10(4), 1794–1805. 10.1021/pr101065j

67. Cox, J., Hein, M. Y., Luber, C. A., Paron, I., Nagaraj, N., & Mann, M. (2014). Accurate Proteome-wide Label-free Quantification by Delayed Normalization and Maximal Peptide Ratio Extraction, Termed MaxLFQ. Molecular & Cellular Proteomics, 13(9), 2513–2526. 10.1074/mcp.M113.031591

68. Schwanhäusser, B., Busse, D., Li, N., Dittmar, G., Schuchhardt, J., Wolf, J., Chen, W., & Selbach, M. (2011). Global quantification of mammalian gene expression control. Nature, 473(7347), 337–342. 10.1038/nature10098

69. Chen Meng. (2022). Fast analyzing, interpreting and sharing quantitative omics data using omicsViewer (1.1.3). Zenodo.

70. Perez-Riverol, Y., Bai, J., Bandla, C., García-Seisdedos, D., Hewapathirana, S., Kamatchinathan, S., Kundu, D. J., Prakash, A., Frericks-Zipper, A., Eisenacher, M., Walzer, M., Wang, S., Brazma, A., & Vizcaíno, J. A. (2022). The PRIDE database resources in 2022: a hub for mass spectrometry-based proteomics evidences. Nucleic Acids Research, 50(D1), D543–D552. 10.1093/nar/gkab1038

