## Supplementary Figures and Tables for "Awakening of the RuMP cycle for partial methylotrophy in the thermophile *Parageobacillus thermoglucosidasius*"

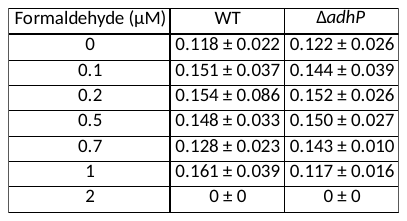

Table S1: Growth rates (h^-1^) of *P. thermoglucosidasius* WT or ∆*adhP* strains growing under different concentrations of formaldehyde.

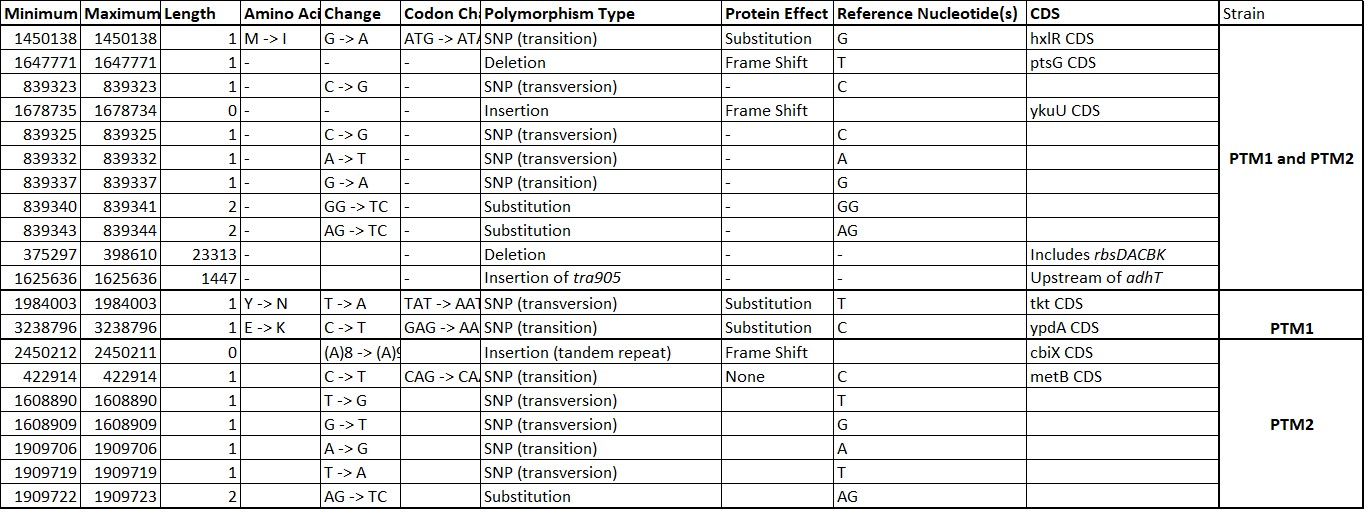

Table S2: All mutations found in the evolved partially methylotrophic strains PTM1 and PTM2.

| M9 minimal medium | Trace elements for M9: | M9 relaxing media: | M9 selective media |
| --- | --- | --- | --- |
| 50 mM 𝑁𝑎_2_𝐻𝑃𝑂_4_ | 134 μM 𝐸𝐷𝑇𝐴 | M9 minimal medium | M9 minimal medium |
| 20 mM 𝐾𝐻_2_𝑃𝑂_4_ | 31 μM 𝐹𝑒𝐶𝑙_2_ | 10 mM Glucose | 10 mM Glucose |
| 1 mM 𝑁𝑎𝐶𝑙 | 6,2 μM 𝑍𝑛𝐶𝑙_2_ | 1 mM Isoleucine | 1 mM Isoleucine |
| 20 mM 𝑁𝐻_4_𝐶𝑙 | 0,76 μM 𝐶𝑢𝐶𝑙_2_ | 1 mM Methionine | 1 mM Threonine |
| 2 mM 𝑀𝑔𝑆𝑂_4_ | 0,42 μM 𝐶𝑜𝐶𝑙_2_ | 1 mM Threonine | 0,25 mM DAP |
| 100 μM 𝐶𝑎𝐶𝑙_2_ | 1,62 μM 𝐻_3_𝐵𝑂_3_ | 0,25 mM DAP |  |
| +trace elements | 0,081 μM 𝑀𝑛𝐶𝑙_2_ | 50 μM 𝑀𝑛𝐶𝑙2 |  |

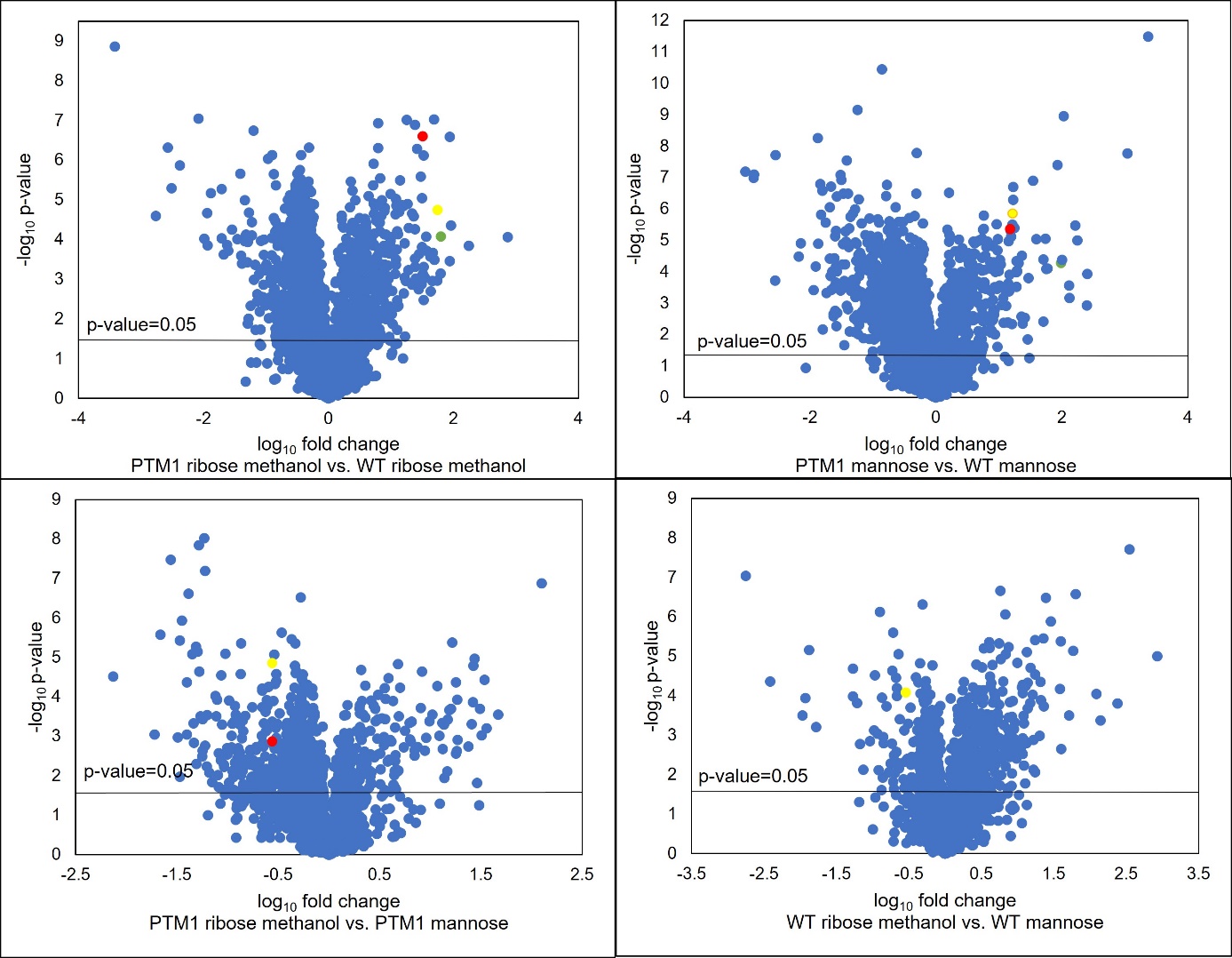
Table S3: Composition of media used for the in vivo assays comprising the *E. coli* formaldehyde biosensor strains.

Figure S1: Volcano plots depicting the abundance difference of all detected proteins and their corresponding t-test p-value. Crucial enzymes for methylotrophy are highlighted in different colors: AdhT: green, Hps (HxlA): yellow, Phi (HxlB): red.

| Name | Sequence |
| --- | --- |
| pJS28-part1-GG-F | tggtctcaatggtcatagctgtttcctgtgtg |
| pJS28-part1-GG-R | tggtctcatagaccatggagagaaaagaaaatcg |
| pJS28-part2-GG-F | tggtctcatctaacttttccactttttgtcttgtcc |
| pJS28-part2-GG-R | tggtctcagcgcggtatcattgcagc |
| pJS28-part3-GG-F | tggtctcagcgcgacccacgctcaccgg |
| pJS28-part3-GG-R | tggtctcagcatgcctgcaggtcgac |
| pJS15A_BsaI_F | tggtctcatcatagctgtttcctgtgtgaa |
| pJS15A_BsaI_R | tggtctcagcatgcctgcaggtc |
| Pbgl_GG_F | tggtctcaatgccgataaacgcgaagaaggtg |
| Pbgl_GG_R | tggtctcatcgaatcactccttatctagaatgcaacctcctttatgttcg |
| TCas9_part1_GG_F | tggtctcctcgaatgaagtataaaatcggtcttgatatc |
| TCas9_part1_GG_R | aggtctcatggtgagccgtactttttg |
| TCas9_part2_GG_F | aggtctcaaccagtctccatccatatcgaactg |
| TCas9_part2_GG_R | tggtctcaccatgattacgccaagcttc |
| USHR_adhP_gib_R | aaggaaagtctggggatcctaccgccgacatcgt |
| DSHR_adhP_gib_F | ggcatgcctgcaggtcgactacgtgttaattgaaggagagacc |
| USHR_rpe_gib_F | aaggaaagtctggggatcctggtgacatttgaaatcccatataagga |
| DSHR_rpe_gib_R | ggcatgcctgcaggtcgactgaatcttcacaaagttcgtattttagtgg |
| Protospacer_rpe_F | aaatgcccatccatcacatcgacgtataacggtatccattttaagaataatccatttttc |
| Protospacer_rpe_R | accgttatacgtcgatgtgatggatgggcatttgtcatagttcccctgagattatcg |
| Protospacer_adhP_F | tcttcattaggcagaccgacaacgtataacggtatccattttaagaataatccatttttc |
| Protospacer_adhP_R | acgttgtcggtctgcctaatgaagagtcatagttcccctgagattatcgc |
| rpe-KO-check2-F | gtataaaacataccgtttgaccggac |
| rpe-KO-check2-R | gcaaataaacgcacgtggaaag |
| adhP-KO-check2-F | gctggcatataaagtggcg |
| adhP-KO-check2-R | cgtcggaatatgtgtcaaacg |
| PZASS_adhT_F | aagaatgcatcatcaccatcaccacaaaactgcgaaaattgttcagc |
| PZASS_adhT_R | ttaaaatttagtgattacgttaaaccctttcg |
| PZASS_backbone_F | gtttaacgtaatcactaaattttaataagctagcgcggcc |
| PZASS_backbone_R | gtggtgatggtgatgatgc |

Table S4: List of primers used.
